# The TRRAP transcription cofactor represses interferon-stimulated genes in colorectal cancer cells

**DOI:** 10.1101/2021.04.13.439699

**Authors:** Dylane Detilleux, Peggy Raynaud, Bérengère Pradet-Balade, Dominique Helmlinger

**Affiliations:** CRBM, University of Montpellier, CNRS, Montpellier, France

**Keywords:** transcription, chromatin, chaperones, PIKKs, interferon, colorectal cancer

## Abstract

Transcription is essential for cells to respond to signaling cues and involves factors with multiple distinct activities. One such factor, TRRAP, functions as part of two large complexes, SAGA and TIP60, which have crucial roles during transcription activation. Structurally, TRRAP belongs to the PIKK family but is the only member classified as a pseudokinase. Recent studies established that a dedicated HSP90 co-chaperone, the TTT complex, is essential for PIKK stabilization and activity. Here, using endogenous auxin- inducible degron alleles, we show that the TTT subunit TELO2 promotes TRRAP assembly into SAGA and TIP60 in human colorectal cancer cells (CRC). Transcriptomic analysis revealed that TELO2 contributes to TRRAP regulatory roles in CRC cells, most notably of MYC target genes. Surprisingly, TELO2 and TRRAP depletion also induced the expression of type I interferon genes. Using a combination of nascent RNA, antibody-targeted chromatin profiling (CUT&RUN) and kinetic analyses, we show that TRRAP directly represses the expression of IRF9, which is a master regulator of interferon stimulated genes. We have therefore uncovered a new, unexpected transcriptional repressor role for TRRAP, which may contribute to its tumorigenic activity.

## INTRODUCTION

Transcriptional regulation is crucial for cells to adapt to external changes, for example during development or to maintain homeostasis. A critical step in gene expression is the initiation of transcription, which is controlled by both *cis*- and *trans*-regulatory mechanisms. These include the coordinated activities of large, multimeric complexes that modify histones or remodel nucleosomes at promoters. These complexes often function as co-activators, bridging DNA-bound transcription factors to the general transcription machinery. By interacting with many transcription factors, co-activators integrate *cis*-regulatory information from multiple inputs and have important roles in establishing specific gene expression programs. The evolutionary conserved SAGA and TIP60 complexes are two paradigmatic examples of such multifunctional co-activator complexes (1, 2). SAGA carries histone acetyltransferase (HAT) and de-ubiquitination (DUB) activities, which preferentially target histone H3 and H2B, respectively. Similar to the general transcription factor TFIID, with which it shares five core TAF subunits, SAGA delivers TBP to specific promoters, stimulating pre-initiation complex (PIC) assembly. TIP60 also carries HAT activity and, through the EP400 subunit, catalyzes deposition of the H2A.Z variant. TIP60 acetylates histone H4 and H2A, as well as the histone variant H2A.Z.

SAGA and TIP60 share one component, named TRRAP in mammals or Tra1 in yeast (3–6). The primary role of Tra1/TRRAP is to recruit SAGA and TIP60 to specific promoters upon binding of an activator. TRRAP was initially discovered as a co-activator for the c-MYC and E2Fs transcription factors and is essential for their oncogenic activities (7). Further work demonstrated that many additional activators require Tra1 or TRRAP to stimulate transcription initiation (8–16). Work in yeast demonstrated that Tra1 physically interacts with the transactivation domain of activators *in vivo* (17–21). Accordingly, genetic studies showed that TRRAP activates the expression of genes involved in a number of important processes. In mammals, these include cell cycle progression (22), mitotic checkpoints (23), the maintenance of a pool of stem or progenitor cells (24–28), and the regulation of cellular differentiation (29). However, the direct regulatory roles of Tra1/TRRAP have been challenging to study, because TRRAP is essential for cell proliferation and early embryonic development in mice (22). Phenotypic analyses have thus relied on partial depletion and conditional knock-out strategies, which are limited by the slow kinetics and irreversibility of the disruption.

TRRAP is an evolutionary conserved member of an atypical family of kinases, called phosphoinositide 3 kinase-related kinases (PIKK), with which it shares a characteristic domain architecture. However, TRRAP lacks all catalytic residues and is therefore the sole pseudokinase of this family (reviewed in (30)). Active PIKKs are implicated in diverse processes. ATM, ATR, and DNA-PKcs control DNA repair and telomere homeostasis, mTOR modulates cell growth, proliferation, and survival in response to metabolic inputs, and SMG-1 mediates nonsense-mediated mRNA decay (30, 31). Elegant studies have demonstrated that PIKKs require a dedicated HSP90 co-chaperone, the triple T complex (TTT), for their stabilization and incorporation into active complexes (32–37). TTT was initially discovered in the fission yeast *Schizosaccharomyces pombe* and is composed of three conserved, specific subunits, TELO2, TTI1, and TTI2 (34, 38, 39). Biochemical evidence suggests a model by which TTT targets the pleiotropic HSP90 chaperone to PIKKs specifically. Mechanistically, TTT recruits HSP90 through the phosphorylation-dependent interaction of TELO2 with the R2TP complex, formed by RPAP3, PIH1D1 and the AAA+ ATPases RUVBL1 and RUVBL2 (40–42). Functional studies in various organisms have implicated TTT, particularly TELO2, in PIKK signaling in response to DNA damage or metabolic stress (32, 33, 35, 36, 43–46). More recent work showed that TTT itself can respond to signaling cues, such as nutrient levels, to modulate PIKK levels, localization, or substrate binding (46–49). In contrast, the effect of TTT on the incorporation of the TRRAP pseudokinase into the SAGA or TIP60 complexes and on their transcription regulatory roles remains poorly characterized, despite evidence that TTT interacts with and stabilizes TRRAP in mammalian cells (32, 35, 36, 45). We recently showed that, in fission yeast, TTT promotes Tra1 stabilization and complex assembly (50). Interestingly, however, *S. pombe* apparently lacks orthologs of the R2TP-specific subunits RPAP3 and PIH1D1 (51), suggesting that the mechanism of PIKK complex assembly may differ between species. In addition, little is known about how the SAGA and TIP60 complexes are assembled in mammals, which chaperones are required, and whether TRRAP is incorporated into each complex by similar or distinct mechanisms.

Here we characterized the role of TTT in gene expression and its contribution to TRRAP regulatory activities. Using CRISPR-Cas9 genome editing, we fused an auxin-inducible degron (AID) to the endogenous TELO2 or TRRAP proteins in human HCT116 colorectal cancer cells (CRC). Biochemical and functional analyses indicated that TELO2 controls TRRAP activity in several ways. First, TELO2 promotes the incorporation of TRRAP into both SAGA and TIP60 complexes. Second, TELO2 regulates the expression of a large fraction of TRRAP-dependent genes. Most genes were previously annotated as targets of c-MYC and E2Fs, suggesting that TTT has an important role in sustaining the activities of these oncogenic transcription factors in CRC cells. Unexpectedly, we also found that both TTT and TRRAP inhibit the expression of type I interferon-stimulated genes (ISGs). Taking advantage of the rapid kinetics and reversibility of the degron allele, we demonstrated that TRRAP is a new, direct transcriptional repressor of the *IRF7* and *IRF9* genes, which encode master regulators of ISG induction. To conclude, our work shows that TRRAP, although a pseudokinase, shares a dedicated chaperone machinery with its cognate kinases for its stability and function. Furthermore, we have identified an unexpected repressive role of TRRAP in the transcription regulation of a subset of genes important for innate immunity and resistance to chemotherapy in CRC cells.

## RESULTS

### Acute and rapid depletion of TELO2 and TRRAP using an auxin-inducible degron

Previous work reported that the TTT components TELO2, TTI1, and TTI2 stabilize TRRAP levels at steady-state (32, 35, 36, 45). We therefore sought to determine whether and how TTT contributes to TRRAP-dependent gene regulation. For this aim, we first asked if TTT promotes the incorporation of TRRAP into the SAGA and TIP60 coactivator complexes.

Our initial attempts to knockdown TTT components in HCT116 CRC cells using RNA interference (RNAi) produced inconclusive results. We reasoned that, to observe the effects of a chaperone on its clients, we need an inducible depletion strategy that targets protein levels directly. Such systems allow assaying phenotypes rapidly after protein depletion and facilitate the ordering of mechanistic events. In addition, a conditional approach was dictated by the observations that both TELO2 and TRRAP are essential for early embryonic development and cell proliferation (22, 32).

For this, we constructed HCT116 cell lines in which an auxin-inducible degron (AID) sequence is integrated at the endogenous *TELO2* or *TRRAP* loci. The AID system mediates proteasomal-dependent degradation of AID-tagged proteins through auxin-mediated interaction with the F-box protein transport inhibitor response 1 (TIR1) (52, 53). We first generated an HCT116 cell line that stably expresses *Oryza sativa* (OsTIR1). One cell line, which we will refer to as parental control (Control), was then used for CRISPR-Cas9- mediated bi-allelic integration of an AID sequence encoding a full-length IAA17-derived degron (Supplemental Figure 1A,B). We also included a YFP fluorescent tag to allow clone selection. We integrated the AID-YFP tag at the 3’end of *TELO2*. In contrast, we targeted the 5’ end of *TRRAP* because its C-terminal FATC domain is critical for function in yeast (54, 55). We also inserted repeated HA epitopes and a 2A peptide (P2A) between the YFP and HA-AID sequences to cleave off the YFP tag during translation, in order to lower the risks of affecting TRRAP function with a long fusion sequence. After transfection, sorting, clonal dilution and amplification, we amplified about 60 clones that showed normal proliferation rates (Supplemental Figure 1C,D). We then screened for a complete size shift of TELO2 and TRRAP by Western blot, using antibodies against the endogenous proteins (Supplemental Figure 1E,F). For each tagged protein, three distinct homozygous clones were isolated and further characterized.

We found that auxin addition induces a rapid and robust depletion of either TELO2^AID^ or ^AID^TRRAP, which levels become undetectable after four hours of treatment (Figure 1A,B). Depletion of either TELO2 or TRRAP progressively decreases the proliferation rate of HCT116 cells (Figure 1C,D and Supplemental Figure 1C,D), consistent with previous results in mice (22, 32). As expected, prolonged TELO2 depletion reduces the steady-state levels of the TRRAP, ATM, ATR, and mTOR proteins and impairs mTORC1 activity, as illustrated with p70-S6K phosphorylation (Supplemental Figure 1E). Similarly, prolonged TRRAP depletion reduces the expression of the Cyclin A2 (*CCNA2*) gene (Supplemental Figure 1F), which is a known target of TRRAP (25, 26, 56). Finally, we verified that cell proliferation and PIKK levels are not affected upon treating the parental, TIR1-expressing control cell line with either auxin or its vehicle (Figure 1C,D and Supplemental Figure 1C-F). In conclusion for this part, we constructed new tools for manipulating TELO2 and TRRAP endogenous levels in a human cell line.

**Figure 1:**
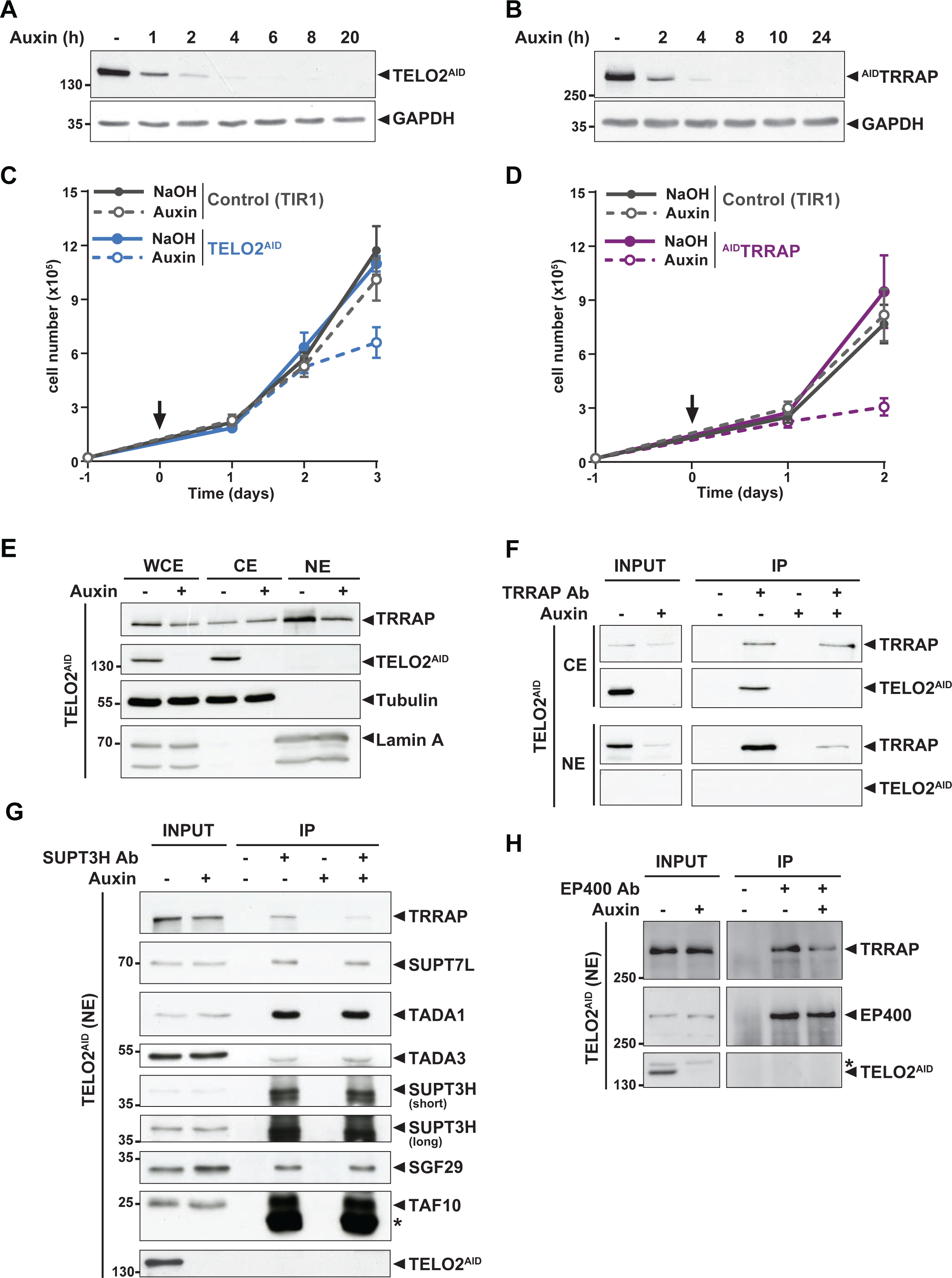
**TELO2 promotes TRRAP assembly into the SAGA and TIP60 complexes.** (A,B) Western-blot analyses of total extracts from TELO2^AID^ (A) or ^AID^TRRAP (B) cell lines harvested at different time points after auxin addition, as indicated (hours). Blots were probed with an anti-TELO2 antibody (A), an anti-HA antibody to detect the HA-AID-TRRAP fusion protein (B), and an anti-GAPDH antibody to control for equal loading. (C,D) Proliferation rates of parental TIR1 (Control, grey lines), TELO2^AID^ (C, blue lines), and ^AID^TRRAP cells (D, purple lines). Cells were seeded one day before treatment (arrow) with either NaOH (solid lines) or auxin (dashed lines) and counted using trypan blue at the indicated time points. Each value represents average cell counts from four independent experiments, overlaid with the standard deviation (SD). (E) Immunoblotting of TRRAP and TELO2^AID^ in whole cell (WCE), cytoplasmic (CE) and nuclear (NE) extracts of TELO2^AID^ cells treated with auxin (+) or NaOH (-) for 48 hours. Tubulin and Lamin A were used as cytoplasmic and nuclear markers, respectively. (F) TRRAP immunoprecipitation (Ab +) from cytoplasmic (CE) and nuclear (NE) fractions of TELO2^AID^ cells treated with auxin (+) or NaOH (-) for 48 hours. TRRAP and TELO2^AID^ were revealed by immunoblotting a fraction (2%) of each extract (INPUT) and the entire immunopurified eluate (IP). (G,H) SUPT3H or EP400 immunoprecipitation (Ab +) from nuclear extracts of TELO2^AID^ cells treated with auxin (+) or NaOH (-) for 24 hours. SUPT3H (G) and EP400 (H), TRRAP, and each indicated SAGA subunits were revealed by immunoblotting a fraction (2%) of the extract (INPUT) and the entire immunopurified eluate (IP). Short and long indicates various exposure times. The star (*) symbol labels antibody light chain contamination (G) or nonspecific detection (H). (F-H) Control IPs (Ab -) were performed using beads only. Data are representative of three independent experiments.

### TELO2 promotes the assembly of TRRAP into the SAGA and TIP60 complexes

We first tested whether TELO2 stabilizes TRRAP at chromatin, where it functions. Cellular fractionation followed by Western blot analysis showed that endogenous TELO2 is detected in the cytoplasm and not in the nucleus (Figure 1E). We observed a similar pattern of the YFP fluorescence signal of TELO2-AID cells using live microscopy (data not shown). In contrast, TRRAP is predominantly nuclear (Figure 1E), consistent with its function and published immunofluorescence microscopy staining (25, 29). Nonetheless, TRRAP can also be observed in the cytoplasmic fraction (Figure 1E). Immunoprecipitation of TRRAP from each compartment showed that TELO2 interacts specifically with the cytoplasmic fraction of TRRAP, but not the nuclear fraction (Figure 1F). We noticed that TELO2 depletion primarily affects nuclear TRRAP levels (Figure 1E,F), suggesting that TELO2 binds newly synthesized TRRAP in the cytoplasm to promote its assembly into SAGA and TIP60, which would then be imported into the nucleus.

We therefore examined the effect of TELO2 on TRRAP incorporation into each complex. For this, we treated TELO2^AID^ cells with auxin for only 24 hours, a time point when TRRAP levels remain unchanged (input in Figure 1G,H). Immunoprecipitation of the SAGA-specific subunit SUPT3H showed a reduced interaction with TRRAP upon TELO2 depletion (Figure 1G). In contrast, TELO2 depletion did not affect the interaction between SUPT3H and the SAGA subunits SUPT7L, TADA1, TADA3, SGF29, and TAF10 (Figure 1G). Similarly, we observed a reduced interaction between TRRAP and the TIP60-specific component EP400 upon TELO2 depletion (Figure 1H). Overall, we conclude that TELO2 promotes TRRAP stability, nuclear localization, and incorporation into the SAGA and TIP60 complexes. These processes are likely tightly coupled with each other, such that unassembled TRRAP is rapidly degraded and not imported into the nucleus.

### SAGA regulates unfolded protein response genes independently of TRRAP

Interestingly, this analysis also revealed that reducing the levels of TRRAP within SAGA does not affect the interaction between SUPT3H, the bait, and all the other SAGA subunits tested (Figure 1G). This observation suggests that TRRAP, despite being about 420 kDa and the largest subunit of SAGA, might be dispensable for SAGA overall integrity and function, as demonstrated in yeast (50, 55). To explore this possibility further, we tested the role of TRRAP in the expression of SAGA-dependent genes during the unfolded protein response (UPR). For this, we monitored UPR gene expression, which requires SAGA for induction in response to endoplasmic reticulum (ER) stress in human cells (57, 58). As expected, we found that two such genes, *CHOP* and *HERPUD*, are strongly induced upon treating HCT116 cells with thapsigargin (Supplemental Figure 2). Consistent with previous work, their induction decreases about two-fold upon RNAi-mediated knockdown of the SAGA core subunit SUPT20H, as compared with control siRNA transfections. In contrast, despite robust TRRAP depletion (Figure 1B), we observed a robust induction of both *CHOP* and *HERPUD* in ^AID^TRRAP cells, irrespective of auxin treatment (Supplemental Figure 2). Taken together, these results suggest that TRRAP does not have a major contribution to SAGA integrity and function at these genes. Thus, despite its large size and absence of catalytic activity, TRRAP is presumably not a core subunit of SAGA in human cells, consistent with recent structural studies revealing its peripheral localization within the complex (59–61).

### TELO2 and TRRAP have overlapping regulatory roles in gene expression

We next sought to characterize the role of TTT in gene expression and its contribution to TRRAP regulatory roles. For this, we performed RNA sequencing (RNA-seq) analyses of TELO2- or TRRAP-depleted cells. Based on the growth curves of three distinct clones of each genotype (Supplemental Figure 1C,D), we treated TELO2^AID^ and ^AID^TRRAP cells for 48 and 24 hours, respectively. No obvious growth defects were observed at these time points, limiting confounding effects from proliferation arrest. Visualization of aligned reads immediately identified several transcripts which levels changed in response to TELO2 and TRRAP depletion. For example, expression of *MCIDAS*, a direct target of TRRAP in multiciliated cells (29), decreases upon TELO2 and TRRAP depletion, while expression of the keratin family gene *KRT20* increases (Supplemental Figure 3A,B).

Differential expression analyses revealed both similarities and differences between the transcriptomes of TELO2- and TRRAP-depleted cells. Using a 1% false discovery rate (FDR) cut-off, we identified 2,188 transcripts which levels are regulated by TRRAP, whereas only 391 transcripts depend on TELO2 (Supplemental Tables 1 and 2). Comparing these two data sets, we found a positive correlation between the gene expression changes caused by TELO2 and TRRAP depletion (Pearson’s correlation coefficient *r* = 0.66) (Figure 2A). Indeed, a majority of genes which expression decreases upon TELO2 depletion (135/168) are downregulated in TRRAP-depleted cells (Figure 2B). Likewise, almost half of the genes which expression increases upon TELO2 depletion (106/223) are upregulated in TRRAP-depleted cells (Figure 2B). Reverse transcription followed by quantitative PCR (RT-qPCR) analyses confirmed these findings for a few selected genes. We observed that TELO2 and TRRAP promote the expression of the *CCNA2* and *MYB* genes, which encode the cell cycle regulator Cyclin A2 and the Myb transcription factor, respectively (Supplemental Figure 3E,F). TELO2 and TRRAP also activate two well-characterized MYC targets genes, *GNL3* and the MiR-17-92a-1 Cluster Host Gene *MIR17HG* (Supplemental Figure 3E,F) (62, 63). Conversely, both TELO2 and TRRAP inhibit the expression of *KRT20*, a marker of differentiated epithelial cells (Supplemental Figure 3G,H). Overall, these observations indicate that TELO2 and TRRAP regulate an overlapping set of genes in HCT116 cells and support our conclusion that TELO2 contributes to TRRAP stability and assembly into SAGA and TIP60 (Figure 1G,H).

**Figure 2:**
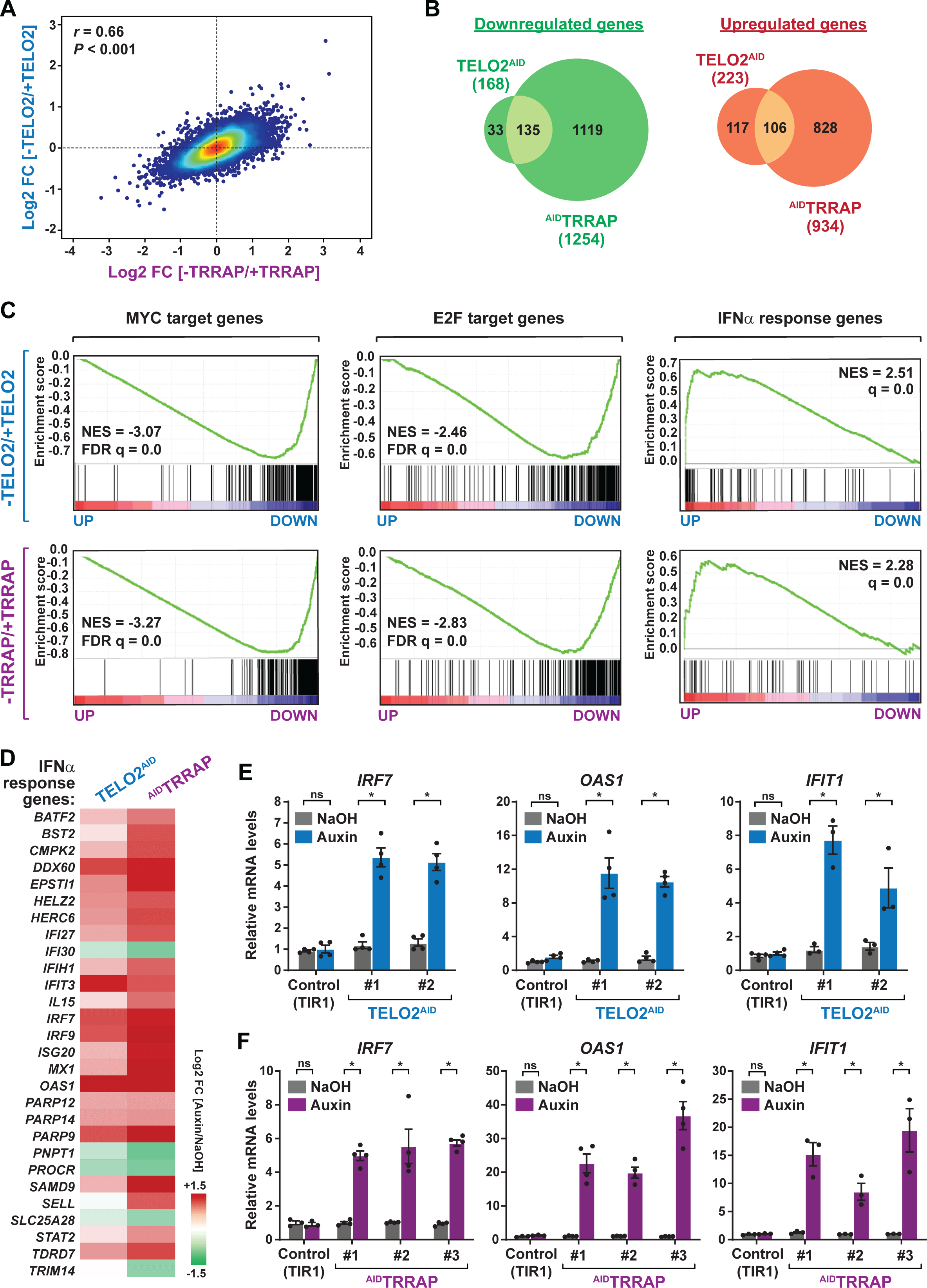
**TELO2 and TRRAP regulate an overlapping set of genes and repress type I interferon stimulated genes.** (A) Density scatter plot comparing gene expression changes between TELO2 and TRRAP- depleted cells. The Pearson correlation coefficient and corresponding *P* value are indicated. (B) Venn diagrams showing the overlap between genes differentially expressed upon TELO2 and TRRAP depletion (FDR = 1%). The number of transcripts which levels decrease and increase are shown separately, as indicated. The number of genes in each set is indicated. (C) Gene set enrichment analysis (GSEA) showing the most strongly enriched hallmarks in the ranked transcriptome profiles of TELO2- (upper panels) and TRRAP-depleted cells (lower panels). Green lines represent enrichment profiles. NES: normalized enrichment score. Each hit from the hallmark gene set is represented by a vertical black bar, positioned on the ranked transcriptome profile with color-coded fold change values. MYC- and E2F- target genes are enriched in the set of genes that are downregulated upon TELO2 and TRRAP depletion, whereas genes from the type I interferon response hallmark are enriched in the set of upregulated genes. (D) Heat map representation of the expression of IFN α-responsive genes (97 genes, hallmark systematic name: M5911) in TELO2 and TRRAP-depleted cells. The Log2 ratio between auxin- and NaOH-treated cells for each transcript is represented using a sequential color scale. All data are from RNA-seq experiments performed in three distinct TELO2^AID^ and ^AID^TRRAP clones treated with auxin or NaOH for 48h or 24h, respectively. (E, F) Quantitative RT-PCR analysis of three ISGs, *IRF7*, *OAS1*, and *IFIT1*, following the depletion of TELO2 (E) and TRRAP (F). mRNAs levels were measured in two TELO2^AID^ clones treated with auxin or NaOH for 48h (E), and in three ^AID^TRRAP clones treated with auxin or NaOH for 24h (F). The corresponding parental control, TIR1-expressing cell lines were also analyzed and treated identically. Each value represents mean mRNA levels from at least three independent experiments, overlaid with individual data points and error bars showing the SD. *PPIB* served as a control for normalization across samples. Values from one NaOH-treated parental control replicate were set to 1, allowing comparisons across culture conditions and replicates. Statistical significance was determined by two-way ANOVA followed by Bonferroni’s multiple comparison tests. *P ≤ 0.01.

We then performed an ontology analysis of genes controlled by TELO2 and TRRAP. We used the Gene Set Enrichment Analysis (GSEA) method and the hallmark gene sets from the Molecular Signatures Database (MSigDB) as a reference (64, 65). We found that the set of genes downregulated upon TRRAP depletion is mostly enriched for MYC or E2Fs target genes (Figure 2C, lower panels). This observation confirms that TRRAP is a co- activator of the c-MYC and E2Fs transcription factors in HCT116 cells, similar to its role in other mammalian cell types. Remarkably, the set of genes activated by TELO2 was also enriched for MYC or E2Fs targets, with comparable enrichment scores (Figure 2C, upper panels). Therefore, TELO2 is a novel regulator of the c-MYC and E2Fs transcriptional factors in HCT116 CRC cells, presumably through its role in promoting TRRAP incorporation into the SAGA and TIP60 co-activator complexes.

Despite these similarities, we also observed differences between each transcriptome profile. First, differentially expressed genes are quantitatively more affected upon TRRAP depletion than upon TELO2 depletion (Figure 2A and Supplemental Figure 3C,D). Second, TRRAP regulates a higher number of genes than TELO2 (Figure 2B and Supplemental Figure 3C,D). Specifically, most genes that are either downregulated (1119/1254) or upregulated (828/934) upon TRRAP depletion remain unchanged in absence of TELO2 (Figure 2B). These differences likely result from the incomplete depletion of TRRAP at this time point of auxin treatment in TELO2^AID^ cells (Figure 1E,F and Supplemental Figure 1E). Finally, a TORC1 signaling hallmark was specifically enriched in the gene set regulated by TELO2, consistent with its well-characterized roles in TORC1 assembly (Supplemental Figure 4A) (34, 36).

### TELO2 and TRRAP repress type I interferon-stimulated genes

Unexpectedly, our ontology analysis also revealed that both TELO2 and TRRAP regulate the expression of genes induced by type I interferons (IFN-I). Using an adjusted *P*-value threshold of 0.05, we found that nearly one third of the 97 interferon-stimulated genes (ISGs) (28/97) from the IFNα response hallmark is differentially expressed in absence of TELO2 and TRRAP (Figure 2C, right panels). However, in contrast to MYC and E2Fs targets, TELO2 and TRRAP repress the expression of most ISGs (23/28), whereas they activate only a few ISGs (5/28) (Figure 2D). Accordingly, the ISG response is the most highly enriched category in the set of genes repressed by TELO2 (NES=2.51) and TRRAP (NES=2.28) (Figure 2C). To strengthen this observation, we performed RT-qPCR analyses of selected ISGs. We measured the expression of *IRF7*, which encodes a transcription factor acting as a master regulator of IFN-I production and signaling, and of two downstream effectors of this pathway, 2’-5’-Oligoadenylate Synthetase 1 *OAS1* and Interferon Induced Protein With Tetratricopeptide Repeats 1 *IFIT1* (66). All three genes exhibit increased mRNA levels upon depletion of either TELO2 or TRRAP (Figure 2E,F). As compared to TELO2, we observed that TRRAP depletion causes a stronger induction of ISGs (Figure 2D-F).

We next verified that auxin itself does not trigger an IFN-I response. Previous work showed that auxin activates genes regulated by the aryl hydrocarbon receptor (AHR), which is a transcription factor responding to diverse chemicals (67). Analysis of the *cis*-regulatory features enriched in the set of genes repressed by TELO2 or TRRAP identified several genes with AHR-binding motifs in their promoters, confirming this finding in HCT116 cells (Supplemental Figure 4B). However, we found that none of the TELO2- or TRRAP-regulated ISGs have AHR-binding motifs within 10 kb of the transcription start sites, except *IFIT3*. Accordingly, filtering out genes with AHR-responsive motifs did not change the results from gene set enrichment analyses. Finally, we verified that auxin treatment of parental HCT116 cells, expressing TIR1 only, does not affect *IRF7*, *OAS1*, and *IFIT1* expression (Figure 2E,F). Altogether, these results show that ISGs are specifically induced in response to TELO2 and TRRAP depletion, indicating that TELO2 and TRRAP repress ISG expression in unstimulated HCT116 CRC cancer cells.

### TRRAP depletion does not activate an innate immune response

TRRAP is best known for its activating role during transcription initiation. Our observation that several ISGs are induced upon TRRAP depletion prompted us to characterize this phenotype further. ISG expression is typically induced in response to pathogens. Infection can trigger various pathogen recognition receptor (PRR) signaling pathways, which converge on the TBK1-mediated phosphorylation and activation of the IRF3 and IRF7 transcription factors (reviewed in (68–70)). IRF3 and IRF7 stimulate the expression of type I IFNs, particularly the interferon-α and –β cytokines, which are then secreted to activate autocrine or paracrine IFN-I signaling through the JAK/STAT pathway. Phosphorylation of STAT1 and STAT2 promotes the recruitment of IRF9 to form the heterotrimeric transcription factor complex ISGF3, which activates the transcription of ISGs to elicit host defense (Figure 3A). Importantly, the *IRF7*, *IRF9*, *STAT1*, and *STAT2* genes are themselves transcriptionally induced by ISGF3 to establish a positive feedback regulatory loop.

**Figure 3:**
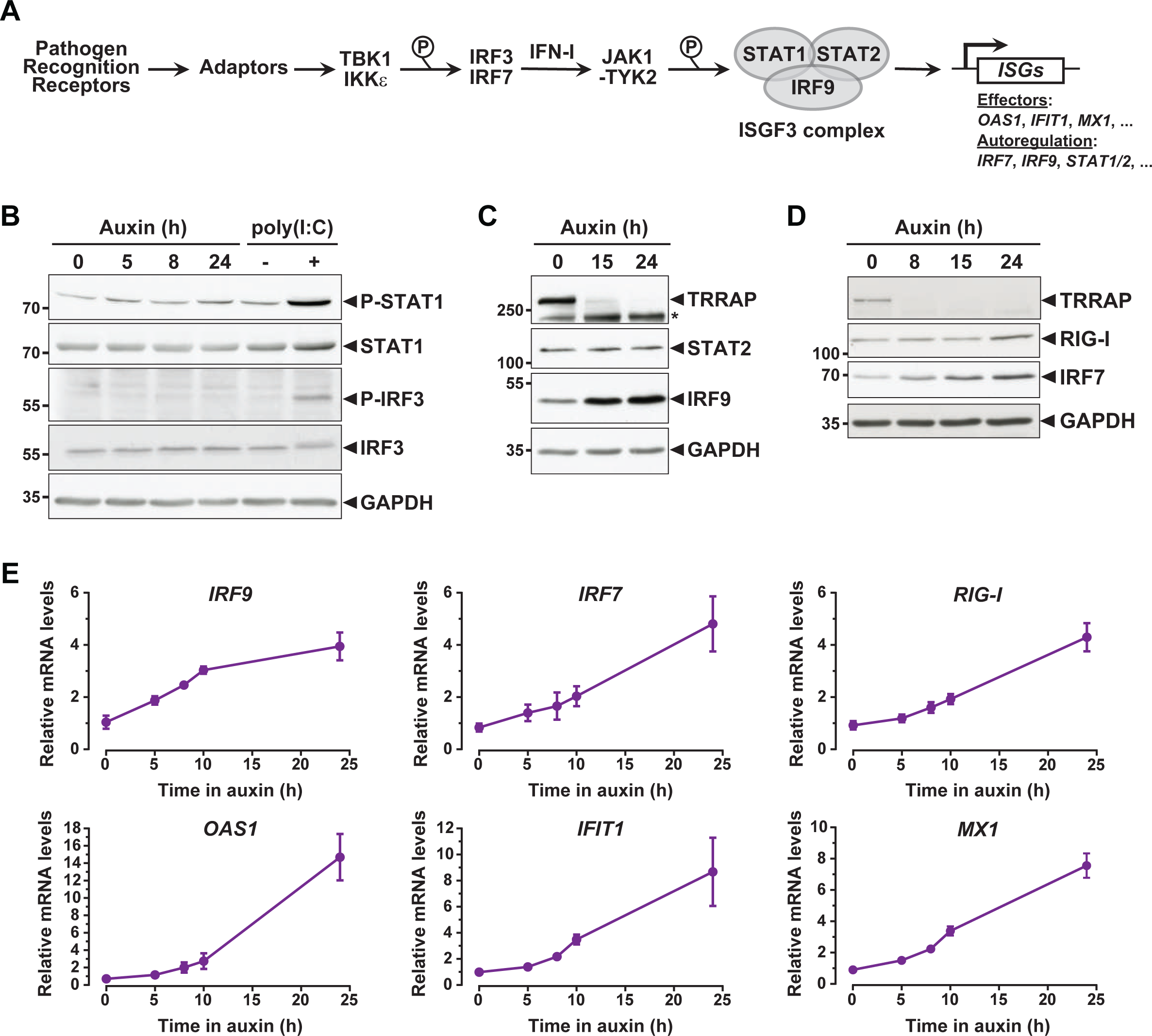
**ISG induction in absence of an innate immune response.** (A) Schematic representation of the innate immune and IFN-I signaling pathways. (B) Western blot analyses of phosphorylated and total STAT1 and IRF3 levels in extracts from ^AID^TRRAP cells treated either with NaOH for 24h and auxin for various time points, or transfected with polyI:C, as indicated. (C) Western blot analyses of TRRAP, STAT2, and IRF9 levels in extracts from ^AID^TRRAP cells treated with NaOH for 24h or auxin for various time points, as indicated. The star (*) symbol labels nonspecific detection. (D) Western blot analyses of TRRAP, RIG-I, and IRF7 levels in extracts from ^AID^TRRAP cells treated with NaOH for 24h or auxin for various time points, as indicated. (E) RT-qPCR analysis of *IRF9*, *IRF7, RIG-I, OAS1, IFIT1*, and *MX1* mRNA levels in ^AID^TRRAP cells over a time course of auxin treatment. RNAs were extracted from cells treated with auxin and harvested at various time points, as indicated. Each value represents mean mRNA levels from at least three independent experiments with distinct ^AID^TRRAP clones, overlaid with error bars showing the SD for each time point. *PPIB* served as a control for normalization across samples. Values from one untreated replicate were set to 1, allowing comparisons across culture conditions and replicates.

To understand how TRRAP regulates ISG expression, we first examined the phosphorylation status of IRF3 and STAT1 in TRRAP-depleted cells, as proxies for the activation of the PRR and IFN-I pathways, respectively. Western blot analyses showed no detectable increase of IRF3 Ser396 phosphorylation and STAT1 Ser727 phosphorylation upon TRRAP depletion (Figure 3B). IRF3 and STAT1 phosphorylation levels remain unchanged both at early and late time points of TRRAP depletion. We noted that IRF3 and STAT1 protein levels did not change, although RNA-seq showed a modest 1.7-fold increase of STAT1 mRNA levels (Supplemental Table 2). As a control, we treated ^AID^TRRAP cells with a double-stranded RNA mimic, polyinosinic acid:polycytidylic acid (poly(I:C)), which is a potent IFN-I inducer (71). As expected, poly(I:C) induced IRF3 and STAT1 phosphorylation, validating our experimental conditions and indicating that a functional innate immune response can be triggered in ^AID^TRRAP HCT116 cell lines. Therefore, TRRAP depletion induces ISG expression without activating the PRR and IFN-I signaling pathways and presumably acts downstream of the IRF3 and IRF7 transcription regulators.

### Kinetic analysis of ISG regulation by TRRAP

While performing this analysis, we noticed that different innate immune response factors accumulate with apparently distinct kinetics in TRRAP-depleted cells (Figure 3C,D). Specifically, the IRF7 and IRF9 transcription factors accumulate earlier than the double-stranded (ds) RNA sensor DDX58 (RIG-I). We thus took advantage of the kinetics of the degron system to better characterize the dynamics of ISG expression upon TRRAP depletion, in an attempt to order phenotypic events. We followed the expression of representative ISGs over a time course of auxin treatment in ^AID^TRRAP cells. Quantitative RT-PCR analyses showed that the relative mRNA levels of *IRF9* increase rapidly after TRRAP depletion. Indeed, expression of *IRF9* is about two-fold higher already 5 hours after auxin addition (Figure 3E), which is concomitant with the loss of a detectable TRRAP signal by Western blot (Figure 1B). However, *IRF9* mRNA levels do not increase noticeably beyond 10 hours of TRRAP depletion and reach a plateau around four-fold. In contrast, *IRF7* mRNA levels gradually increase over 24 hours of TRRAP depletion and accumulate with a small delay as compared to *IRF9* (Figure 3E). Importantly, Western blot analyses showed the same trend. IRF7 progressively accumulates over the time-course of TRRAP depletion, whereas IRF9 begins to level off at an earlier time point, around 15 hours (Figure 3C,D). Finally, we found that the expression of other ISGF3 targets, including *RIG-I*, *OAS1*, *IFIT1*, and *MX1* gradually increases upon TRRAP depletion, similar to *IRF7* (Figure 3E). Likewise, Western blotting showed that RIG-I upregulation is detectable only 24 hours after auxin addition (Figure 3D). Altogether, the progressive accumulation of ISGs indicates that this phenotype appears early after the loss of TRRAP, rather than at late time points. Furthermore, this kinetic analysis indicates that the IRF9 transcription factor is induced earlier than its target genes, which include *IRF7*.

### TRRAP represses a specific subset of ISGs

We next sought to determine how TRRAP modulates ISG expression without activating an innate immune response or the IFN-I signaling pathway. Accumulating evidence highlights the existence of non-canonical regulatory mechanisms of ISG transcription. Notably, high levels of IRF9 alone are sufficient to trigger ISG transcription in HCT116 cells (72). Indeed, IRF9 and STAT2 can form STAT1-independent complexes that drive specific transcriptional programs (73). Here, our RNA-seq, quantitative RT-PCR and Western blot analyses indicate that IRF9 levels increase about four-fold (Supplemental Table 2, Figure 3C,E). In contrast, STAT1 and STAT2 levels do not change (Figure 3C). Thus, TRRAP depletion induces a progressive accumulation of IRF9, but neither activation nor overexpression of its partners STAT1 and STAT2.

Work in both normal and cancer cells demonstrated that IRF9 can assemble with unphosphorylated STAT1 and STAT2 to form the unphosphorylated ISGF3 (U-ISGF3) complex. U-ISGF3 induces the transcription of a subset of 29 ISGs and sustains their prolonged, constitutive expression (74–76). In contrast, the phosphorylated form of ISGF3 activates an additional 49 ISGs upon acute IFN-I signaling (Figure 4A). We thus compiled a list of 148 ISGs, combining the ISGs from the MSigDB IFNα response hallmark gene set (65) with the U-ISGF3- and ISGF3-regulated ISGs defined in (74). Of these, 113 ISGs were detected and quantified in our RNA-seq experiments (Supplemental Table 3). Using an adjusted *P*-value threshold of 0.05, we found that TRRAP regulates the mRNA levels of about a third of all ISGs (39/113), mostly negatively (30/39) (Supplemental Figure 5). Interestingly, we noticed that the effect of TRRAP on all 113 ISGs shows a bimodal distribution (grey violin plot, Figure 4B), suggesting that TRRAP regulates a specific subset of ISGs. To compare the effect of TRRAP on U-ISGF3- and ISGF3-regulated genes, we plotted each set separately and observed that TRRAP primarily represses U-ISGF3- dependent genes, while ISGF3-dependent genes remain largely unaffected (compare green and orange violin plots, Figure 4B). Specifically, TRRAP depletion induces the expression of 23 of the 28 U-ISGF3-dependent genes, but only 7 of the 35 ISGF3-dependent genes.

**Figure 4:**
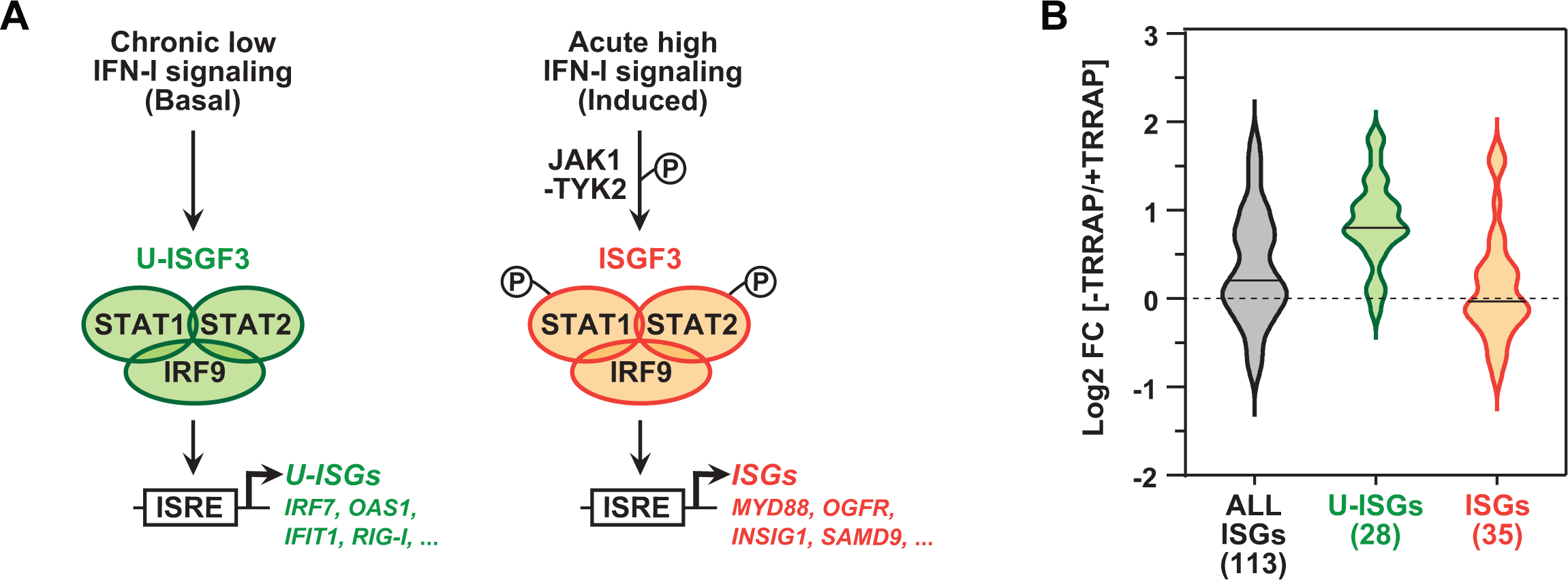
**TRRAP specifically represses U-ISGF3-regulated genes.** (A) Schematic representation of the transcription regulation of distinct subsets of ISGs, depending on whether ISGF3 unphosphorylated, in basal conditions (U-ISGF3, left) or phosphorylated in response to IFN-I signaling (ISGF3, right). (B) Violin plots showing the distribution of gene expression changes induced by TRRAP depletion for all ISGs (grey), for ISGs controlled by U-ISGF3 (green), and for ISGs controlled by ISGF3 (orange). Log2 fold changes (Log2[FC]) were calculated as the Log2 of the ratio of the expression value of each gene between NaOH- (+TRRAP) and auxin-treated (-TRRAP) ^AID^TRRAP cells. The number of genes in each category is indicated.

Finally, we determined whether specific transcription factor binding sites are enriched in the set of 39 ISGs regulated by TRRAP, using the i-*cis*Target and Pscan tools (77, 78). This analysis identified the interferon-stimulated response element (ISRE) as the most enriched motif in the promoter regions of TRRAP-regulated ISGs (Supplemental Figure 5). ISREs are specifically recognized by IRF9 homo- or heterodimers, as well as the ISGF3 complex (70), suggesting that IRF9 upregulation is sufficient to induce ISG expression in TRRAP-depleted cells, independently of IFN-I signaling. IRF7-binding sites were the second most enriched motifs in the set of TRRAP-regulated ISGs. In agreement with this observation, IRF7 protein levels progressively accumulate upon TRRAP depletion (Figure 3D), consistent with the 2.6- fold increase observed by RNA-seq (Supplemental Table 2).

In conclusion, TRRAP inhibits the expression of a specific subset of ISGs in HCT116 cells. These ISGs are coregulated by U-ISGF3, the unphosphorylated form of the IRF9- STAT1-STAT2 complex, and can be induced independently of activation of the innate immune and IFN-I signaling pathways.

### TRRAP directly represses *IRF7* and *IRF9* mRNA synthesis

Our results so far suggest that TRRAP is a negative regulator of IRF7 and IRF9 which, once overexpressed, would induce a specific subset of ISGs. We therefore focused on the regulation of the IRF7 and IRF9 transcription factors. Indeed, TRRAP has well-established roles in activation of transcription, but not repression. We therefore sought to determine if TRRAP directly represses the transcription of the *IRF7* and *IRF9* genes.

Our expression analyses were limited to measurements of steady-state mRNA levels, which correspond to the net balance between RNA synthesis and decay. In order to measure the effect of TRRAP on *IRF9* and *IRF7* mRNA synthesis, we performed metabolic labeling of newly synthesized RNAs in TRRAP-depleted cells. For this, ^AID^TRRAP cells were treated with auxin, followed by a short incubation with 4-thiouridine (4sU). 4sU-labelled RNAs were then purified and quantified by conventional quantitative RT-PCR. To verify the efficiency of labeling and purification, we amplified intronic regions of each gene. We found that the levels of newly synthesized *IRF7* and *IRF9* pre-mRNAs increase about 1.8- and 1.6- fold in absence of TRRAP, respectively (Figure 5A). Upon TRRAP depletion, we observed an 8-fold increase in *OAS1* transcription, as expected for a target of the IRF7 and IRF9 activators. We conclude that TRRAP inhibits *IRF7* and *IRF9* mRNA synthesis, suggesting that TRRAP functions as a transcriptional repressor for these genes.

**Figure 5:**
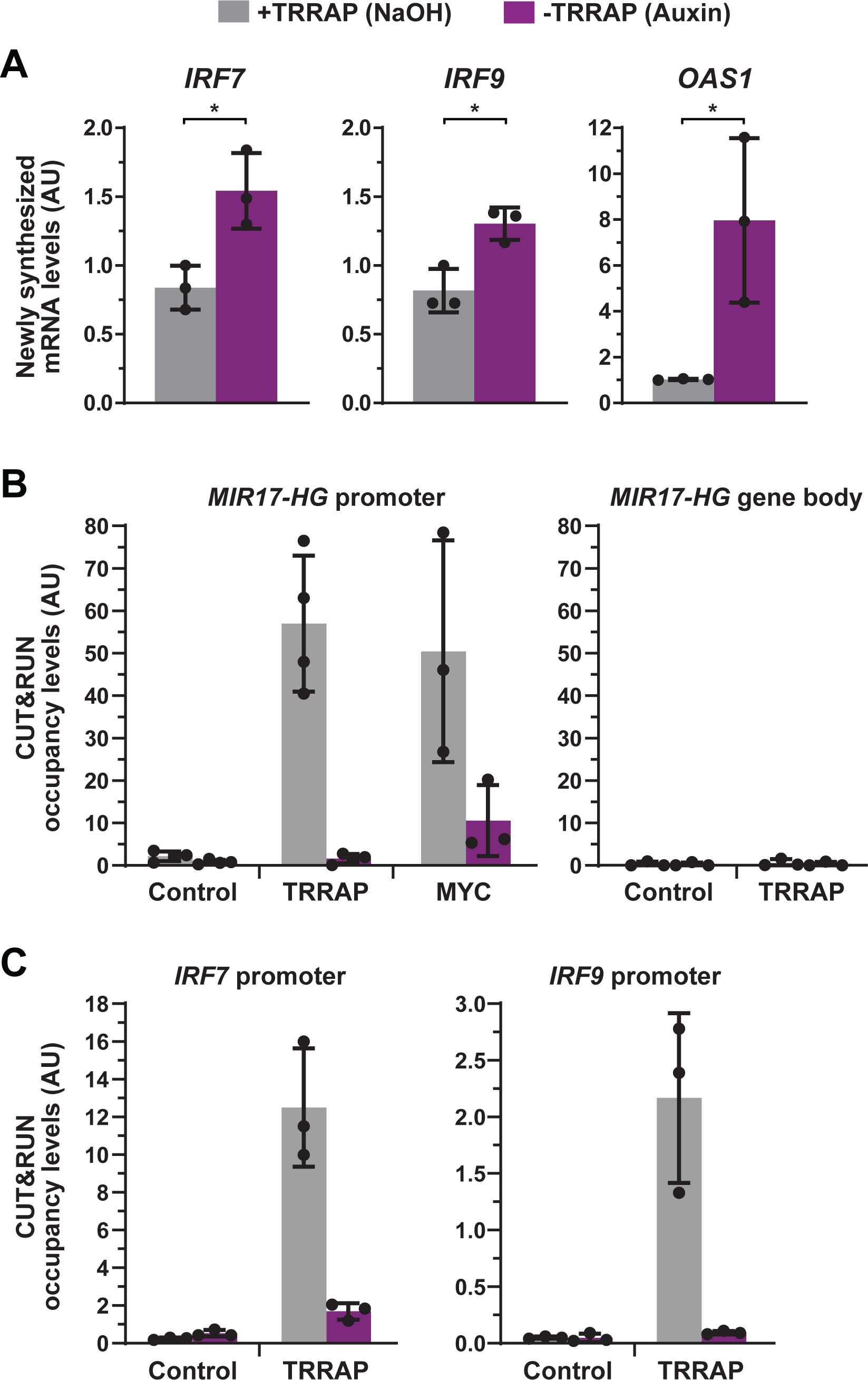
**TRRAP directly represses the transcription of *IRF7* and *IRF9*.** (A) RT-qPCR analysis of newly synthesized *IRF7*, *IRF9,* and *OAS1*. ^AID^TRRAP cells with either NaOH (grey) or auxin (purple) for 10 hours, prior to a 20-minute incubation with 4sU, extraction, and enrichment of labeled nascent RNAs. Each value represents mean pre- mRNA levels from three independent experiments with distinct ^AID^TRRAP clones, overlaid with individual data points and error bars showing the SD. *PPIB* served as a control for normalization across samples. For all genes, primers amplifying intronic regions were used. Values from one untreated replicate were set to 1, allowing comparisons across culture conditions and replicates. Statistical significance was determined by unpaired, two-tailed Student’s *t* tests. *P ≤ 0.05. (B) CUT&RUN-qPCR analysis of TRRAP and MYC occupancy at the *MIR17-HG* locus in ^AID^TRRAP cells with either NaOH (grey) or auxin (purple) for 12 hours. CUT&RUN were performed using rabbit IgGs (control), anti-HA (TRRAP) and MYC antibodies. Cleaved DNA fragments were released, extracted, and analyzed by qPCR using oligonucleotides that amplify either the promoter or the gene body of *MIR17-HG*. Each column represents the mean value of at least three independent experiments with distinct ^AID^TRRAP clones, overlaid with individual data points and error bars showing the SD. (C) CUT&RUN-qPCR analysis of TRRAP occupancy at the *IRF7* and *IRF9* promoter regions in ^AID^TRRAP cells with either NaOH (grey) or auxin (purple) for 12 hours.

In order to distinguish between direct and indirect regulatory effects, we next asked whether TRRAP binds to the promoter regions of the *IRF7* and *IRF9* genes. Although chromatin immunoprecipitation (ChIP) is typically used to identify genomic regions occupied by transcription regulators, this procedure suffers from poor signal-to-noise ratio, particularly for factors that do not bind DNA directly, such as TRRAP and TRRAP-containing complexes. To overcome this problem, we implemented a novel profiling strategy, called Cleavage Under Targets and Release Using Nuclease (CUT&RUN) (79). This technique is based on targeted endogenous cleavage of chromatin by micrococcal nuclease (MNase), which is directed to a protein using a specific antibody. Here, after release and purification, we analyzed cleaved DNA fragments using conventional quantitative PCR. Overall, CUT&RUN avoids formaldehyde crosslinking, which can yield misleading results and mask epitopes, and generates very little noise because undigested chromatin and genomic DNA are not extracted.

To validate the entire CUT&RUN-qPCR procedure, we first assessed TRRAP chromatin occupancy at a well-characterized MYC-bound locus. We reasoned that TRRAP binding would mirror that of MYC because they interact both physically and functionally in many cell lines. Accordingly, our RNA-seq analysis shows that TRRAP activates MYC target genes in HCT116 cells (Figure 2C). We selected the *MIR17-HG* gene, which encodes an oncomir precursor, is a prominent target of MYC in cancer cells (63), and is activated by TRRAP in HCT116 cells (Supplemental Figure 3F). Analysis of published MYC CUT&RUN-seq data confirmed that it binds to a large region of the *MIR17-HG* promoter in human K562 chronic myeloid leukemia cells (80). Accordingly, we found that MYC binds to the promoter of *MIR17-HG* in ^AID^TRRAP HCT116 cells (Figure 5B), validating the CUT&RUN-qPCR procedure.

To measure TRRAP binding, we repeated the CUT&RUN-qPCR procedure, using an HA antibody that recognizes the repeated HA epitopes fused to the N-terminal end of ^AID^TRRAP (Figure 1B and Supplemental Figure 1B). Our analysis shows that this strategy is remarkably efficient to profile TRRAP chromatin occupancy (Figure 5B). Indeed, we observed a strong footprint of TRRAP at the promoter region of *MIR17-HG*. Comparison with several negative controls confirmed the robustness and specificity of this observation. First, TRRAP binding at *MIR17HG* is about 60-fold above that observed using rabbit IgGs. Second, enrichment at the promoter is specific because background signals are detected downstream, in the gene body. Third, auxin-mediated TRRAP depletion reduces the HA signal to background levels (Figure 5B). Interestingly, TRRAP depletion also reduces the binding of MYC to *MIR17-HG*, suggesting that TRRAP stabilizes MYC interaction with its promoters. Finally, we examined the binding profile of TRRAP at the *IRF7* and *IRF9* genes. CUT&RUN-qPCR analyses revealed robust and specific binding of TRRAP to the proximal promoter regions of both *IRF7* and *IRF9*, as compared to control IgGs and TRRAP-depleted conditions (Figure 5C).

In summary, combining nascent RT-qPCR analyses and an innovative chromatin profiling technique, we show that TRRAP binds to the *IRF7* and *IRF9* promoters and represses their transcription. We therefore conclude that, unexpectedly, TRRAP is a direct transcriptional repressor of these master regulators of ISG expression in HCT116 cells. Additionally, we validated the CUT&RUN technique for profiling the occupancy of a component of large chromatin-modifying and -remodeling complexes in human cells.

### Dynamics of ISG regulation by TRRAP

To further support a direct role for TRRAP in repressing *IRF7* and *IRF9*, we studied the dynamics of their expression and of TRRAP promoter recruitment, taking advantage of the reversibility of the AID system (52). For this, after complete auxin-induced TRRAP depletion, we transferred ^AID^TRRAP cells to culture medium lacking auxin for various time points (Figure 6A). Western blot analyses confirmed that, following initial depletion, TRRAP progressively re-accumulates within a few hours after washing out auxin from the media (Figure 6B).

**Figure 6:**
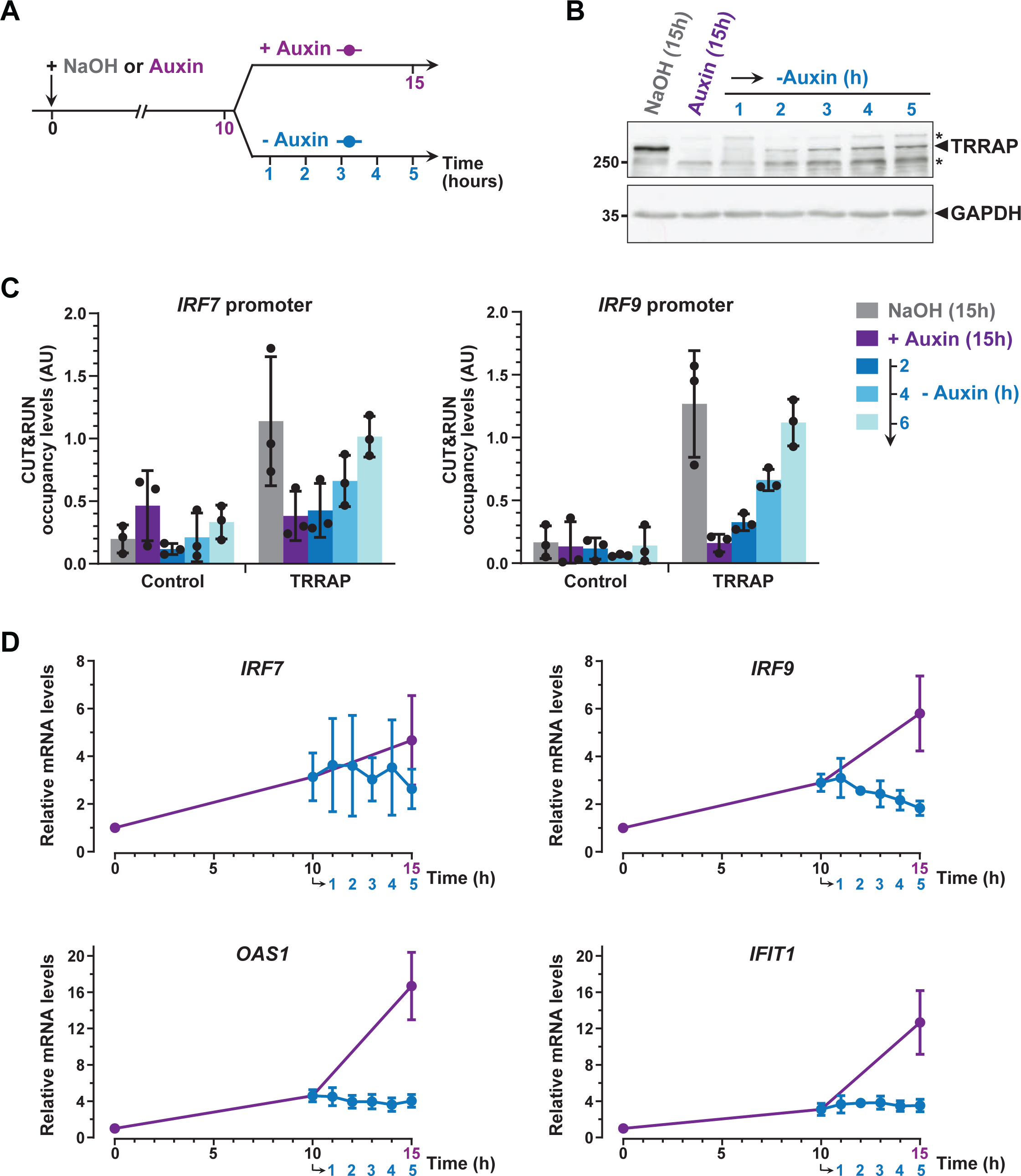
**Dynamics of ISG regulation by TRRAP.** (A) Experimental strategy: ^AID^TRRAP cells were harvested either before auxin addition, or after incubation with NaOH (grey) and auxin for 10 hours (purple), and at various time points up to 5 hours after auxin removal (blue), as indicated. As a control, ^AID^TRRAP cells were also harvested following 15 hours of auxin treatment (purple). (B) Western blot analysis of TRRAP protein levels after auxin addition and subsequent removal, demonstrating the reversibility of TRRAP auxin-mediated degradation. Blots were probed with anti-TRRAP and anti-GAPDH antibodies. The latter was used to control for equal loading. The star (*) symbols label nonspecific detections. (C) CUT&RUN-qPCR analysis of TRRAP-bound DNA extracted following MNase cleavage directed by either an anti-HA antibody (TRRAP) or control IgG (Control). CUT&RUN was performed in ^AID^TRRAP cells that were either with NaOH (grey) or treated with auxin for 15 hours (purple), then washed out of auxin for 2, 4, and 6 hours (blue). qPCR was performed using oligonucleotides that amplify proximal regions of the *IRF7* and *IRF9* promoters, as in Figure 5C. Each column represents the mean value of three distinct ^AID^TRRAP clones, overlaid with individual data points and error bars showing the SD. (D) RT-qPCR analysis of *IRF9*, *IRF7, OAS1,* and *IFIT1* mRNA levels after auxin addition and subsequent removal. RNAs were extracted from cells treated as schematized in (A). Each point represents the mean value of three distinct ^AID^TRRAP clones, overlaid with error bars showing the SD. *PPIB* served as a control for normalization across samples. Values from untreated samples were set to 1, allowing comparisons across culture conditions and replicates.

CUT&RUN followed by qPCR analysis showed that TRRAP binding to *IRF7* and *IRF9* increases upon auxin removal (Figure 6C). Specifically, we observed that TRRAP binding progressively re-appears at both the *IRF7* and *IRF9* promoters, in parallel to TRRAP recovery (Figure 6B,C). Six hours after auxin wash out, we found that TRRAP occupancy is similar to that observed in untreated cells, suggesting that TRRAP binding is highly dynamic. We then sought to correlate TRRAP occupancy profile with its regulatory role in ISG expression. Quantitative RT-PCR of *IRF7* mRNA levels showed no measurable changes, although we noticed a small decrease at later time points of auxin wash out (Figure 6D). In contrast, upon auxin removal, *IRF9* mRNA levels rapidly stop increasing and progressively decrease back to basal levels, as compared with a continuous auxin treatment (Figure 6D). Similarly, TRRAP re-appearance stabilizes the levels of the *OAS1* and *IFIT1* mRNAs, while their expression continues to increase when TRRAP depletion is sustained (Figure 6D).

In conclusion, re-expressing TRRAP leads to a rapid reversion of the ISG induction phenotype. We note that the dynamics of *IRF7* and *IRF9* regulation shows slightly different profiles and suggest a more important role of TRRAP on *IRF9*. Taken together, our kinetic analyses of TRRAP binding and regulatory roles indicate that TRRAP and ISG levels dynamically anti-correlate in HCT116 cells and provide additional evidence for a direct regulatory role for TRRAP in *IRF9* expression.

### Both TIP60 and SAGA contribute to ISG repression

TRRAP regulates transcription as part of either the TIP60 or the SAGA co-activator complex (30). We thus asked whether SAGA and TIP60 contribute to ISG repression in HCT116 cells. Using small-interfering RNAs (siRNAs), we knocked down specific subunits of each complex, targeting their enzymatic activities. As a positive control, we measured ISG expression upon siRNA-mediated knockdown of TRRAP (Figure 7A). Then, for the TIP60 complex, we targeted EP400, an ATPase catalyzing the deposition of histone variant H2A.Z, and the HAT subunit TIP60, which promotes histone H4 acetylation (Figure 7B) (1). For the SAGA complex, we targeted TADA3, a subunit of the HAT module and essential for SAGA HAT activity towards histone H3, and ATXN7L3, which is essential for SAGA DUB activity on histone H2B (Figure 7C) (2). RT-qPCR analyses revealed that *IRF9* and *OAS1* mRNA levels increase upon knockdown of EP400 (Figure 7D). The effect of EP400 knockdown is similar to that observed upon TRRAP knockdown. ATXN7L3 knockdown also induces *IRF9* and *OAS1* expression, although its effect is less pronounced than that of EP400 and TRRAP (Figure 7D). In contrast, we found that downregulating TIP60 has no effect on *IRF9* expression, while it causes a modest increase of *OAS1* levels (Figure 7D). Finally, TADA3 knockdown has no effect on both *IRF9* and *OAS1* expression (Figure 7D). Although differences in the efficiency of each knock-down complicate such comparisons, these analyses indicate that both TRRAP-containing complexes, TIP60 and SAGA, contribute to ISG repression in HCT116 cells, using distinct regulatory activities. Indeed, our findings suggest a model by which ISG repression might depend on H2A.Z deposition by TIP60 and SAGA H2B de-ubiquitylation by SAGA, whereas SAGA and TIP60 HAT activities do not have a major regulatory role in basal conditions.

**Figure 7:**
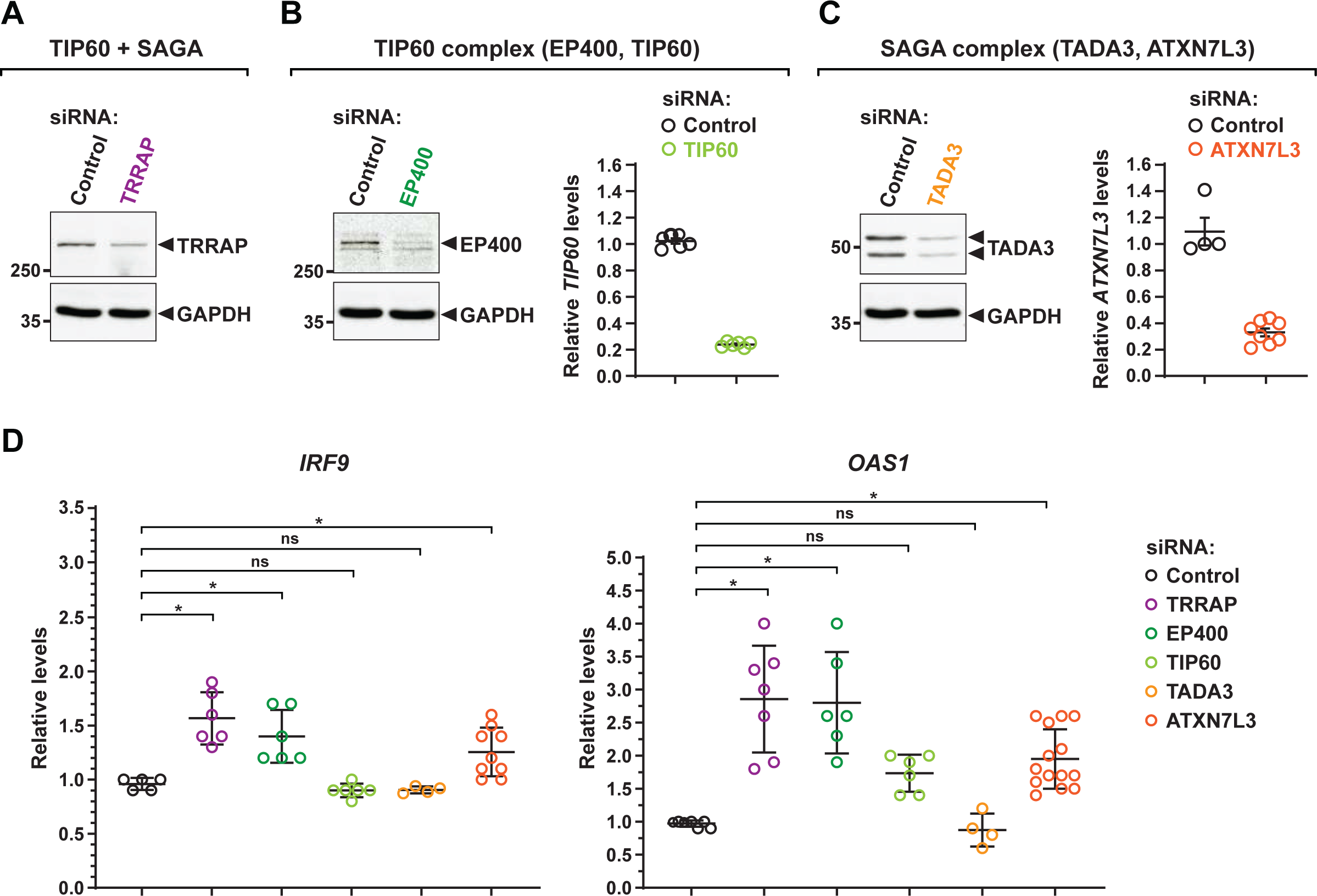
**Contribution of TIP60 and SAGA activities to ISG expression.** (A-C) Western blot and RT-qPCR analyses of siRNA efficiency targeting TRRAP, TIP60- and SAGA-specific subunits. TRRAP (A), EP400 (B), and TADA3 (C) expression was examined using specific antibodies to probe whole cell extracts prepared from HCT116 cells transiently transfected with the corresponding siRNAs, as indicated. An anti-GAPDH antibody was used to control for equal loading across samples. TIP60 (B) and ATXN7L3 (C) expression was examined using RT-qPCR analysis of RNA extracted from cells transiently transfected with the corresponding siRNAs, as indicated. (D) RT-qPCR analysis of *IRF9* (left) and *OAS1* (right) mRNA levels from cells treated as described in (A,B). Each line represents the mean value of at least four independent experiments, overlaid with individual data points and error bars showing the SD. Expression levels were normalized to *PPIB* levels and to cells transfected with a control siRNA targeting Firefly Luciferase (FFL). Values from one siRNA-control sample were set to 1, allowing comparisons across culture conditions and replicates. Statistical significance was determined by one-way ANOVA followed by Tukey’s multiple comparison tests. *P ≤ 0.05; ns: P > 0.05.

## DISCUSSION

TRRAP is a member of the PIKK family of atypical kinases. Genetic and biochemical evidence indicate that the HSP90 cochaperone TTT is an essential regulator of PIKK stability and activity. However, TRRAP lacks catalytic residues and is the sole pseudokinase amongst the PIKK family. Although previous studies showed that TTT interacts with TRRAP and promotes its stability, the contribution of TTT to TRRAP functions remained unknown. Here, using an inducible degron system that allows acute and robust depletion of endogenous proteins, we demonstrate that the TTT subunit TELO2 promotes TRRAP assembly into functional complexes, SAGA and TIP60, and contributes to TRRAP regulatory roles in gene expression. Functional and transcriptomic analyses indicate that both TELO2 and TRRAP promote proliferation of CRC cells and regulate an overlapping set of genes, including MYC targets. Our work also uncovered an unexpected negative role of TRRAP on the expression of a subset of type I interferon stimulated genes. Notably, we accumulated evidence that TRRAP acts as a transcriptional repressor at the promoter of the *IRF9* gene, which encodes a subunit of a trimeric transcription factor complex acting as a master regulator of ISG expression. Our findings suggest that TRRAP and its chaperone TTT might contribute to colorectal tumorigenesis by promoting a specific transcriptional program.

### Chaperone-mediated biogenesis and regulation of the TRRAP pseudokinase

Studies in mammals revealed that the pleiotropic HSP90 chaperone is specifically recruited to PIKKs by a dedicated cochaperone, the TTT complex, to promote their stability and incorporation into active complexes (32–36, 45). In contrast, the effect of TTT on the TRRAP pseudokinase, the only inactive PIKK, is less characterized. Previous work showed that TTT stabilizes TRRAP in human cells (32, 35, 36, 45). Furthermore, the yeast ortholog of TRRAP, Tra1, also requires TTT and Hsp90 for stability, incorporation into the NuA4 and SAGA co-activator complexes, and function in gene expression (50, 81). Here, conditional and rapid depletion of endogenous TELO2 allowed us to better characterize the role of TTT in TRRAP biogenesis and functions in human cells. Our results suggest a model by which TELO2 promotes the cytoplasmic assembly of TRRAP-containing complexes, TIP60 and SAGA, and regulates the expression of a large number of TRRAP-dependent genes. Altogether, these studies indicate that TTT functional roles are conserved between yeast and mammals. Therefore, although Tra1/TRRAP is the sole catalytically inactive member of the PIKK family, it shares a dedicated chaperone machinery with active PIKK kinases for assembly and function. Phylogenetic analyses of PIKK orthologs show that Tra1/TRRAP appeared early in the eukaryotic lineage, concomitantly with all other PIKKs (30). Although catalytic residues diverge, all TRRAP orthologs harbor the distinctive domain architecture of PIKKs, including the highly conserved FAT and kinase domains. Similar to PIKKs, orthologs of TELO2 and TTI1 are found in the genomes of species representative of all major eukaryotic clades (30). We thus propose that the requirement of PIKKs for a dedicated cochaperone explains the selection pressure observed on the sequence and domain organization of Tra1/TRRAP, in the absence of catalytic activity.

Finally, our work indicates that assembly of the NuA4/TIP60 and SAGA complexes requires the HSP90 cochaperone TTT. A recent study showed that human TAF5 and its paralog TAF5L, which are specific to the general transcription factor TFIID and SAGA, respectively, require the CCT chaperonin, for incorporation into pre-assembled modules (82). An alternative mechanism, involving cotranslational interactions between other SAGA subunits, was recently described in yeast and human cells (83, 84). Our study therefore contributes to the emerging concept that dedicated mechanisms and chaperones control the *de novo* assembly of transcription complexes.

### Inhibitory roles of TRRAP in ISG expression

We show here that TELO2 and TRRAP regulate the expression of an overlapping set of genes. Notably, we found that TELO2 is important for the activation of MYC and E2F target genes (Figure 2). In agreement, TRRAP is essential for transcription activation by MYC and E2F and contributes to their oncogenic functions during tumorigenesis (7–10, 85).

Unexpectedly, we found that TELO2 and TRRAP also repress several genes, particularly those mediating the type I interferon (IFN-I) response during innate immunity.

The IFN-I signaling pathway is divided into two branches that define an early and a late response, and involves five major transcription factors. Following detection of “non-self” elements, such as pathogens, PRR signaling leads to activation of the IRF3 and IRF7 transcription factors and production of type I interferons during the early response. Then, these cytokines activate autocrine and paracrine JAK kinase signaling, to induce the formation of the ISGF3 transcription factor complex during the late response. ISGF3 comprises the IRF9, STAT1, and STAT2 transcription factors, which activate several ISGs encoding downstream effectors of IFN-I signaling and innate immunity. ISGF3 also activates *IRF7*, *IRF9*, *STAT1*, and *STAT2* expression to constitute a positive feedback loop (Figure 3A). Our functional analyses indicate that neither the PRR nor the IFN-I signaling pathways are activated upon TRRAP depletion. Rather, we found that TRRAP represses a specific subset of ISGs by directly inhibiting *IRF7* and *IRF9* transcription. First, TRRAP depletion induces ISGs independently of PRR and IFN-I signaling pathway activation, as shown by the lack of IRF3 and STAT1 phosphorylation, respectively (Figure 3B). Second, time course analyses suggest that IRF7 and IRF9 accumulation precedes that of its target genes (Figures 3C-E). Third, TRRAP specifically represses a subset of ISGs, which previous work showed can be induced by unphosphorylated IRF9 overexpression (72, 74–76, 86) (Figure 4). Fourth, TRRAP dynamically binds to the promoters of *IRF7 and IRF9* and represses their transcription (Figures 5A,C and 6). Finally, although both IRF7 and IRF9 likely contribute to ISG induction upon TRRAP depletion, *IRF9* appears as the major target of TRRAP. Indeed, ISGF3 is a well-characterized effector of IFN-I signaling and, as such, functions downstream of TBK1 and IRF7 activation. In addition, IRF9 can activate *IRF7* expression itself, including in HCT116 cells (72). Notably, the rapid kinetics, efficiency, and reversibility of the auxin- inducible degron allowed us to deduce a putative sequence of phenotypic events. We propose that TRRAP depletion relieves *IRF9* repression, which subsequently induces the transcription of *IRF7* and a subset of other ISGs, independently of IFN-I signaling. Nevertheless, we cannot formally exclude that TRRAP activates the transcription of a yet uncharacterized repressor of ISGs. Examination of the list of genes which expression decrease upon TRRAP depletion did not identify an obvious candidate.

How does TRRAP function as a transcriptional repressor for specific ISGs? Our data suggest that both the TIP60 and SAGA complexes are involved. Specifically, we observed that the TIP60 subunit EP400 and the SAGA DUB subunit ATXN7L3 contribute to ISG repression in HCT116 cells. The EP400 ATPase is the catalytic subunit responsible for deposition of the histone variant H2A.Z at +1 nucleosomes. Supporting the involvement of EP400, a proximity labeling screen identified EP400 as a putative interactor of IRF9 (76). Furthermore, H2A.Z is removed during ISGF3-dependent transcription activation and, conversely, H2A.Z silencing induces ISGF3 recruitment to ISG promoters, ISG mRNA expression, and IFN-stimulated antiviral immunity (87). However, we did not detect a reproducible effect of TRRAP on H2A.Z occupancy at the *IRF9* promoter (our unpublished observations). One possibility is that EP400 represses *IRF9* expression by another, unknown mechanism.

Two previous studies showed that SAGA and TIP60 and SAGA restrict the response to innate immunity signaling pathways (88, 89). However, our work suggests that TRRAP functions by a distinct mechanism. One study showed that the SAGA subunit GCN5/PCAF represses IFN-I signaling by inhibiting the TBK1 kinase, which targets IRF3 (88). In contrast, we observed that TRRAP depletion induces ISG expression without IRF3 phosphorylation and activation of IFN signaling. The other study identified TIP60 as a repressor of endogenous retroviral elements (ERVs), which can activate innate immune signaling pathways when depressed (89–91). However, we did not detect activation of an innate immune response in TRRAP-depleted cells. Moreover, both our and their studies show that TIP60 depletion does not induce expression of IRF9 and several target ISGs, suggesting a different mechanism. In contrast, a recent study in *Drosophila* indicates that the Tip60 subunit E(Pc) attenuates expression of components of JAK/STAT signaling (92), which is reminiscent of our findings and suggest that this function of TRRAP might be conserved in other eukaryotic species.

### ISG expression in colorectal cancer

Alterations of TIP60 and SAGA activities have been documented in various cancers, for example as essential cofactors of pro-oncogenic transcription factors (93, 94). The shared subunit TRRAP represents a compelling example because it was identified in a screen for factors that are essential for MYC-dependent transcription and malignant transformation (7). Supporting this conclusion, TRRAP is recurrently overexpressed in CRC patient samples (our analysis of the COADREAD cohort, http://xena.ucsc.edu/) and we show here that TRRAP depletion primarily affects a MYC target signature in HCT116 cells. In addition, our work suggests that TRRAP might contribute to colorectal tumorigenesis by restricting ISG expression levels. Indeed, several studies reported that ISGs are normally repressed in CRC cells, but upregulated upon acquisition of radio- and chemoresistance (74, 95–99). Specifically, gene expression profiling identified a subset of ISGs, defined as the interferon-related DNA damage signature (IRDS), which increases in cancer cells that become resistant to ionizing radiation, topoisomerase inhibitors, or DNA damage. Remarkably, the IRDS is virtually identical to the subset of ISGs induced by unphosphorylated U-ISGF3 and those repressed by TRRAP in HCT116 CRC cells. Therefore, TRRAP might play a new role in colorectal tumorigenesis by repressing a subset of interferon-stimulated genes.

## MATERIALS & METHODS

### Cell culture, reagents and transfection

HCT116 cells (a gift from Dr. Vogelstein, Howard Hughes Medical Institute, Baltimore, MD) were cultured in McCoy’s 5A medium (Sigma) supplemented with 10% (v/v) FBS and 100 U/ml Penicillin/Streptomycin in 5% CO2 at 37°C. For siRNA knockdown, HCT116 cells were seeded in 6-well plates 24 hours before transfection and transfected with 20 nM of siRNA using INTERFERin® (Polyplus Transfections) according to the manufacturer’s protocol. All siRNA sequences used are listed in Supplemental Table S4. Cells were then harvested 48 hours after transfection for RNA isolation and protein analysis. For auxin treatment, 3- indoleacetic acid (IAA) (Sigma, I2886) was dissolved in NaOH 1N and used at a final concentration of 0.5 mM. For interferon pathway activation, 2 µg of polyinosinic-polycytidylic acid (poly(I:C) LMW) (InvivoGen, tlrl-picw) were transfected. For cell viability assay, cells were seeded at 20,000 cells per well in 1 mL in 24-well plates. Cells were counted daily using the Countess^TM^ automated cell counter based on trypan blue exclusion assay (Invitrogen).

### Generation of CRISPR-Cas9-edited cell lines

Single guide RNAs (sgRNAs) were designed using the http://crispr.mit.edu/ website. Then, candidate sgRNAs with the highest scores (indicating fewest potential off-targets) were selected and synthesized. Two complementary oligonucleotides of sgRNAs were annealed and cloned into the BbsI sites of pUC57 vector (Supplemental Table S5). The two different sgRNA plasmids, the pX335-U6-Chimeric_BB-CBh-hSpCas9n(D10A) plasmid (Addgene, #42335) and the donor plasmid containing the AID and YFP cassettes were co-transfected into an HCT116 cell line stably expressing *Os*TIR1-9Myc, using the FuGENE6 transfection reagent (Promega). YFP-positive cells were selected and isolated by fluorescence-activated cell sorting 48 hours after transfection.

### Cell fractionation

Separation of cytoplasmic and nuclear fractions was achieved using the Rapid, Efficient and Practical (REAP) protocol (100). Briefly, culture medium was removed from cell culture dishes and cells were washed twice using ice-cold PBS. 1 ml of PBS was added to each 10 cm dish before cells were scraped and collected in microcentrifuge tubes. Samples were centrifuged (10 seconds at 2000 g) and supernatants were discarded. 1 ml of 0.1% NP40 – PBS1x was added to each pellet with pipetting up and down several times. Small aliquots (150 µL) were set aside as whole cell lysates (WCE). The rest of the samples were centrifuged and supernatants were transferred to new tubes, constituting the cytoplasmic fractions (150 µL) (CE). Pellets were re-suspended in 1 ml 0.1% NP40 and centrifuged to obtain nuclear fractions as pellets (NE). Whole lysates were mixed with 4X Laemmli buffer (ratio 3:1), sonicated and boiled. Cytoplasmic fractions were mixed with 4X Laemmli buffer (ratio 3:1) and boiled. Nuclear fractions were resuspended in 200 µl of 1X Laemmli buffer, sonicated and boiled. For western blot analyses, 20 µL of WCE and CE fractions and 10 µL of NE were loaded.

### Cell lysis and Western blots

Cells from 6-well plates were harvested by trypsinization and lysed in RIPA buffer (20 mM Tris, pH 7.5, 150 mM NaCl, 1% Nonidet-P40, 0.5% sodium deoxycholate, protease inhibitors (cOmplete EDTA-free cocktails tablets, Roche). Cell lysates were subjected to 6% or 10% SDS-PAGE and 40μg of protein were loaded per lane. The gels were transferred for 2 hours onto nitrocellulose membranes (GE Healthcare Biosciences, Pittsburgh, PA). Detection was performed by diluting primary antibodies in TBS supplemented with 0.1% Tween and either 5% BSA or 5% non-fat dry milk. All antibodies used in this study are listed in Supplemental Table S6. After incubation and washing, secondary anti-mouse IgG or anti-rabbit IgG antibodies conjugated to horse-radish-peroxidase (Santa Cruz) were added at a dilution of 1:5,000 in TBS 0.1% Tween 5% non-fat drymilk. Detection was performed using Pierce^TM^ ECL western blotting Substrate (ThermoFisher scientific).

### RT–qPCR analysis

Total RNA was isolated from HCT116 cells using TRIzol reagent (Invitrogen, 15596018), according to the manufacturer’s instructions, followed by a DNAse digestion step using the TURBO DNA-free^TM^ kit (Ambion, AM1907). Total RNA (1 μg) was reverse-transcribed using the SuperScript III reverse transcriptase (Invitrogen, Life Technologies). Real-time quantitative PCR (qPCR) was performed using SYBR Green Master Mix and the LightCycler® 480 instrument (Roche). Relative levels of gene expression were analyzed using the 2ΔΔCt method and compared to the expression of the human housekeeping gene *PPIB*. The cycling conditions comprised an initial denaturation phase at 95°C for 5 minutes, followed by 50 cycles at 95°C for 10 seconds, 60°C for 30 seconds and 72°C for 15 seconds. All oligonucleotides used for amplification are listed in Supplemental Table S5.

### Nascent RT–qPCR analysis

Metabolic labelling of newly transcribed RNAs was performed as described in (101). Briefly, the nucleoside analogue 4-thiouridine (4sU) (Abcam, ab143718) was added to culture medium at a final concentration of 500 μM for a 20 minutes pulse. 4sU was removed by washing cells with ice cold 1x PBS and immediately lysed using TRIzol reagent (Invitrogen, 15596018). Total RNA was extracted according to the manufacturer’s instruction, precipitated, resuspended in 130 μl and sonicated on a Covaris E220 instrument (1 % duty factor, 100 W, 200 cycles per burst, 80 seconds). For purification, fragmented total RNA was incubated for 10 minutes at 60°C and immediately chilled on ice for 2 minutes to melt secondary RNA structures. 4sU-labelled RNA was thiol-specific biotinylated by addition of 200 μg EZ-link HPDP-biotin (ThermoFisher Scientific, 21341), biotinylation buffer (10 mM Hepes-KOH pH 7.5 and 1 mM EDTA) and 20% DMSO (Sigma-Aldrich, D8418) to prevent precipitation of HPDP-biotin. Biotinylation was carried out for 3 hours at 24°C in the dark and with gentle agitations. After incubation, biotin excess was removed by adding an equal volume of chloroform and centrifugation at 16,000 g for 5 minutes at 4°C. RNA was precipitated from the aqueous phase using 0.1 volumes of 5 M NaCl and one volume of 100% isopropanol followed by centrifugation at 16,000 g for 30 minutes at 4°C. After washing with 75% ethanol the RNA pellet was resuspended in 100 μl of RNase-free water and denatured for 10 minutes at 65°C followed by immediate chilling on ice for 5 minutes. RNA samples were incubated with 100 μl of streptavidin-coated μMACS magnetic beads (Miltenyi Biotec, 130-074-101) for 90 minutes at 24°C under gentle agitations. The μMACS columns (Miltenyi Biotec, Cat# 130-074-101) were placed on a MACS MultiStand (Miltenyi Biotec) and equilibrated with washing buffer (100 mM Tris-HCl pH 7.5, 10 mM EDTA, 1 M NaCl, 0.1% Tween20) before applying the samples twice to the columns. The columns were then washed one time with 600 μl, 700 μl, 800 μl, 900 μl and 1 ml washing buffer before eluting the newly synthesized RNA with two washes of 100 μl 0.1M DTT. Purified newly synthesized RNAs were recovered using the RNeasy MinElute Cleanup Kit (Qiagen, 74204), according to the manufacturer’s instruction. 1 μg RNA was reverse-transcribed using the SuperScript III reverse transcriptase (Invitrogen, Life Technologies). Real-time quantitative PCR (qPCR) was performed using SYBR Green Master Mix and the LightCycler® 480 instrument (Roche). Relative levels of gene expression were analyzed using the 2ΔΔCt method and compared to the expression of the human housekeeping gene *PPIB*. The cycling conditions comprised an initial denaturation phase at 95°C for 5 minutes, followed by 50 cycles at 95°C for 10 seconds, 60°C for 30 seconds and 72°C for 15 seconds. All oligonucleotides used for amplification are listed in Supplemental Table S5.

### RNA-seq

Total RNA was extracted using the TRIzol reagent (Invitrogen, 15596018) according to the manufacturer’s instructions, followed by DNAse digestion step using the TURBO DNA-free^TM^ kit (Ambion, AM1907). RNA concentration and quality were analyzed using an Agilent Bioanalyzer 2100 (Agilent Technologies). Total RNA-seq libraries were generated from 1 μg of total RNA using TruSeq Stranded Total RNA LT Sample Prep Kit. High-throughput sequencing of RNA libraries was performed following the standard protocol from Fasteris (www.fasteris.com). All sequencing runs were performed on an Illumina HiSeq 2500.

Transcripts were purified by polyA-tail selection. Stranded dual-indexed cDNA libraries were constructed using the Illumina TruSeq Stranded mRNA Library Prep kit. Library size distribution and concentration were determined using an Agilent Bioanalyzer. 12 libraries were sequenced in one lane of an Illumina HiSeq 4000, with 1 × 50 bp single reads, at Fasteris SA (Plan-les-Ouates, Switzerland). After demultiplexing according to their index barcode, the total number of reads ranged from 23 to 27 million per library. Adapter sequences were trimmed from reads in the Fastq sequence files. Reads were aligned using HISAT2 (102), with strand-specific information (–rnastrandness R) and otherwise default options to human genome assembly GRCh37 (hg19). For all samples, the overall alignment rate was over 80%, including over 90% of reads mapping uniquely to the human genome. Reads were then counted for exon features using htseq-count (103) in union mode (–mode union), reverse stranded (–stranded Reverse), and a minimum alignment quality of 10 (– minaqual 10). For all samples, over 95% of reads were assigned to a feature (–type gene). Variance mean dependence was estimated from count tables and tested for differential expression based on a negative binomial distribution, using DESeq2 (104). Pairwise comparison or one-way analysis of variance were run with a parametric fit and ‘treatment as the source of variation (factor: auxin or NaOH). All computational analyses were run either on R (version 3.6.0), using publicly available packages, or on the Galaxy web platform using the public server at usegalaxy.org (105).

### CUT&RUN

CUT&RUN experiments were implemented based on the procedure described in (79). Briefly, cells were grown up to 80% confluence. Fresh cell cultures were harvested and cells were counted. 250,000 cells were used per sample. Cells were washed and incubated for 10 minutes at room temperature with 10µL of concanavalin A-coated beads. Cells bound to beads were permeabilized and incubated with the appropriate antibody overnight at 4°C. Protein A-MNAse at a final concentration of 700 ng/mL was mixed and incubated with cells at 4°C for 1 hour. pA-MN was activated with 2 µL of 100 mM CaCl_2_ and digestion was performed for 30 minutes at 0°C. The reaction was stopped with 100 µL of stop buffer containing 2 pg/mL of heterologous spike-in DNA. Release of the fragments was achieved at 37°C during 15 minutes. DNA was extracted following using NucleoSpin columns (Macherey-Nagel) and eluted in 30 µL of NE buffer. Extracted DNA was used for qPCR analysis using SYBR Green Master Mix and the LightCycler® 480 instrument (Roche). All oligonucleotides used for amplification are listed in Supplemental Table S5.

### Statistics

Statistical tests were performed using GraphPad Prism. t-Tests were used when comparing two means. One-way or two-way analyses of variance (ANOVA) were performed for comparing more than two means, across one (for example “genotype”) or two distinct variables (for example “genotype” and “treatment”). ANOVAs were followed by Bonferroni or Tukey post hoc pairwise comparisons. Comparisons that are statistically significant (P ≤ 0.05) are marked with star signs (*), whereas those that are statistically not significant (P > 0.05) are labeled n.s.

## Data availability

The raw sequencing data reported in this publication have been deposited in NCBI Gene Expression Omnibus and are accessible through GEO Series accession number GSE171454.

## Supporting information

AllSupplementalFiguresandTables4-6

## ACKNOWLEDGMENTS

We thank Manon Le Goff, Aida Yazbeck, and Boutaina El Kenz for invaluable technical assistance and all members of the Helmlinger laboratory for helpful suggestions and discussions. We are grateful to Véronique Gire, Benjamin Vitre, Daniele Fachinetti, Edouard Bertrand, Didier Devys, and Laszlo Tora for kindly sharing plasmids and cell lines. We are indebted to Véronique Fischer and Steve Henikoff for sharing reagents and helping set up the nascent RNA and CUT&RUN analyses, respectively. We thank Léo Pioger, Emeric Dubois and Samia Guendouz for their help with RNA-seq. D.D. is a recipient of a graduate fellowship from la Ligue Nationale Contre le Cancer. This work was supported by funds from the Fondation ARC (PJA-20131200471) and the INCa grant PLBIO 2016-161.

## REFERENCES

1. Lu, P.Y., Lévesque, N. and Kobor, M.S. (2009) NuA4 and SWR1-C: two chromatin-modifying complexes with overlapping functions and components. Biochem. Cell Biol., 87, 799–815.

2. Helmlinger, D. and Tora, L. (2017) Sharing the SAGA. Trends Biochem. Sci., 42, 850–861.

3. Grant, P.A., Schieltz, D., Pray-Grant, M.G., Yates, J.R. and Workman, J.L. (1998) The ATM-related cofactor Tra1 is a component of the purified SAGA complex. Mol. Cell, 2, 863– 867.

4. Vassilev, A., Yamauchi, J., Kotani, T., Prives, C., Avantaggiati, M.L., Qin, J. and Nakatani, Y. (1998) The 400 kDa subunit of the PCAF histone acetylase complex belongs to the ATM superfamily. Mol. Cell, 2, 869–875.

5. Allard, S., Utley, R.T., Savard, J., Clarke, A., Grant, P., Brandl, C.J., Pillus, L., Workman, J.L. and Côté, J. (1999) NuA4, an essential transcription adaptor/histone H4 acetyltransferase complex containing Esa1p and the ATM-related cofactor Tra1p. EMBO J., 18, 5108–19.

6. Saleh, A., Schieltz, D., Ting, N., McMahon, S.B., Litchfield, D.W., Yates, J.R., Lees-Miller, S.P., Cole, M.D. and Brandl, C.J. (1998) Tra1p is a component of the yeast Ada·Spt transcriptional regulatory complexes. J. Biol. Chem., 273, 26559–26565.

7. McMahon, S.B., Van Buskirk, H.A., Dugan, K.A., Copeland, T.D. and Cole, M.D. (1998) The novel ATM-related protein TRRAP is an essential cofactor for the c-Myc and E2F oncoproteins. Cell, 94, 363–374.

8. Park, J., Kunjibettu, S., McMahon, S.B. and Cole, M.D. (2001) The ATM-related domain of TRRAP is required for histone acetyltransferase recruitment and Myc-dependent oncogenesis. Genes Dev., 15, 1619–1624.

9. Bouchard, C., Dittrich, O., Kiermaier, A., Dohmann, K., Menkel, A., Eilers, M. and Lüscher, B. (2001) Regulation of cyclin D2 gene expression by the Myc/Max/Mad network: Myc-dependent TRRAP recruitment and histone acetylation at the cyclin D2 promoter. Genes Dev., 15, 2042–2047.

10. Lang, S.E., McMahon, S.B., Cole, M.D. and Hearing, P. (2001) E2F Transcriptional Activation Requires TRRAP and GCN5 Cofactors. J. Biol. Chem., 276, 32627–32634.

11. Deleu, L., Shellard, S., Alevizopoulos, K., Amati, B. and Land, H. (2001) Recruitment of trrap required for oncogenic transformation by E1A. Oncogene, 20, 8270–8275.

12. Ard, P.G., Chatterjee, C., Kunjibettu, S., Adside, L.R., Gralinski, L.E. and McMahon, S.B. (2002) Transcriptional Regulation of the mdm2 Oncogene by p53 Requires TRRAP Acetyltransferase Complexes. Mol. Cell. Biol., 22, 5650–5661.

13. Lang, S.E. and Hearing, P. (2003) The adenovirus E1A oncoprotein recruits the cellular TRRAP/GCN5 histone acetyltransferase complex. Oncogene, 22, 2836–2841.

14. Memedula, S. and Belmont, A.S. (2003) Sequential recruitment of HAT and SWI/SNF components to condensed chromatin by VP16. Curr. Biol., 13, 241–246.

15. Knutson, B.A. and Hahn, S. (2011) Domains of Tra1 important for activator recruitment and transcription coactivator functions of SAGA and NuA4 complexes. Mol. Cell. Biol., 31, 818–831.

16. Lin, L., Chamberlain, L., Zhu, L.J. and Green, M.R. (2012) Analysis of Gal4-directed transcription activation using Tra1 mutants selectively defective for interaction with Gal4. Proc. Natl. Acad. Sci. U. S. A., 109, 1997–2002.

17. Brown, C.E., Howe, L., Sousa, K., Alley, S.C., Carrozza, M.J., Tan, S. and Workman, J.L. (2001) Recruitment of HAT complexes by direct activator interactions with the ATM- related Tra1 subunit. Science, 292, 2333–2337.

18. Bhaumik, S.R. and Green, M.R. (2001) SAGA is an essential in vivo target of the yeast acidic activator Gal4p. Genes Dev., 15, 1935–1945.

19. Bhaumik, S.R., Raha, T., Aiello, D.P. and Green, M.R. (2004) In vivo target of a transcriptional activator revealed by fluorescence resonance energy transfer. Genes Dev., 18, 333–343.

20. Fishburn, J., Mohibullah, N. and Hahn, S. (2005) Function of a eukaryotic transcription activator during the transcription cycle. Mol. Cell, 18, 369–78.

21. Reeves, W.M. and Hahn, S. (2005) Targets of the Gal4 Transcription Activator in Functional Transcription Complexes. Mol. Cell. Biol., 25, 9092–9102.

22. Herceg, Z., Hulla, W., Gell, D., Cuenin, C., Lleonart, M., Jackson, S. and Wang, Z.Q. (2001) Disruption of Trrap causes early embryonic lethality and defects in cell cycle progression. Nat. Genet., 29, 206–211.

23. Li, H., Cuenin, C., Murr, R., Wang, Z.-Q.Q. and Herceg, Z. (2004) HAT cofactor Trrap regulates the mitotic checkpoint by modulation of Mad1 and Mad2 expression. EMBO J., 23, 4824–4834.

24. Loizou, J.I., Oser, G., Shukla, V., Sawan, C., Murr, R., Wang, Z.-Q.Q., Trumpp, A. and Herceg, Z. (2009) Histone acetyltransferase cofactor Trrap is essential for maintaining the hematopoietic stem/progenitor cell pool. J. Immunol., 183, 6422–6431.

25. Wurdak, H., Zhu, S., Romero, A., Lorger, M., Watson, J., Chiang, C.-Y. yuan, Zhang, J., Natu, V.S., Lairson, L.L., Walker, J.R., et al. (2010) An RNAi screen identifies TRRAP as a regulator of brain tumor-initiating cell differentiation. Cell Stem Cell, 6, 37–47.

26. Tapias, A., Zhou, Z.W., Shi, Y., Chong, Z., Wang, P., Groth, M., Platzer, M., Huttner, W., Herceg, Z., Yang, Y.G., et al. (2014) Trrap-dependent histone acetylation specifically regulates cell-cycle gene transcription to control neural progenitor fate decisions. Cell Stem Cell, 14, 632–643.

27. Tauc, H.M., Tasdogan, A., Meyer, P. and Pandur, P. (2017) Nipped-A regulates intestinal stem cell proliferation in Drosophila. Development, 144, 612–623.

28. Sawan, C., Hernandez-Vargas, H., Murr, R., Lopez, F., Vaissière, T., Ghantous, A.Y., Cuenin, C., Imbert, J., Wang, Z.-Q.Q., Ren, B., et al. (2013) Histone acetyltransferase cofactor Trrap maintains self-renewal and restricts differentiation of embryonic stem cells. Stem Cells, 31, 979–991.

29. Wang, Z., Plasschaert, L.W., Aryal, S., Renaud, N.A., Yang, Z., Choo-Wing, R., Pessotti, A.D., Kirkpatrick, N.D., Cochran, N.R., Carbone, W., et al. (2018) TRRAP is a central regulator of human multiciliated cell formation. J. Cell Biol., 217, 1941–1955.

30. Elías-Villalobos, A., Fort, P. and Helmlinger, D. (2019) New insights into the evolutionary conservation of the sole PIKK pseudokinase Tra1/TRRAP. Biochem. Soc. Trans., 47, 1597–1608.

31. Lempiäinen, H. and Halazonetis, T.D. (2009) Emerging common themes in regulation of PIKKs and PI3Ks. EMBO J., 28, 3067–73.

32. Takai, H., Wang, R.C., Takai, K.K., Yang, H. and de Lange, T. (2007) Tel2 Regulates the Stability of PI3K-Related Protein Kinases. Cell, 131, 1248–1259.

33. Anderson, C.M., Korkin, D., Smith, D.L., Makovets, S., Seidel, J.J., Sali, A. and Blackburn, E.H. (2008) Tel2 mediates activation and localization of ATM/Tel1 kinase to a double-strand break. Genes Dev., 22, 854–859.

34. Takai, H., Xie, Y., de Lange, T. and Pavletich, N.P. (2010) Tel2 structure and function in the Hsp90-dependent maturation of mTOR and ATR complexes. Genes Dev., 24, 2019–2030.

35. Hurov, K.E., Cotta-Ramusino, C. and Elledge, S.J. (2010) A genetic screen identifies the Triple T complex required for DNA damage signaling and ATM and ATR stability. Genes Dev., 24, 1939–1950.

36. Kaizuka, T., Hara, T., Oshiro, N., Kikkawa, U., Yonezawa, K., Takehana, K., Iemura, S.-I., Natsume, T. and Mizushima, N. (2010) Tti1 and Tel2 are critical factors in mammalian target of rapamycin complex assembly. J. Biol. Chem., 285, 20109–20116.

37. Izumi, N., Yamashita, A., Iwamatsu, A., Kurata, R., Nakamura, H., Saari, B., Hirano, H., Anderson, P. and Ohno, S. (2010) AAA+ proteins RUVBL1 and RUVBL2 coordinate PIKK activity and function in nonsense-mediated mRNA decay. Sci. Signal., 3, ra27.

38. Hayashi, T., Hatanaka, M., Nagao, K., Nakaseko, Y., Kanoh, J., Kokubu, A., Ebe, M. and Yanagida, M. (2007) Rapamycin sensitivity of the Schizosaccharomyces pombe tor2 mutant and organization of two highly phosphorylated TOR complexes by specific and common subunits. Genes to cells, 12, 1357–1370.

39. Shevchenko, A., Roguev, A., Schaft, D., Buchanan, L., Habermann, B., Sakalar, C., Thomas, H., Krogan, N.J., Shevchenko, A. and Stewart, A.F. (2008) Chromatin Central: towards the comparative proteome by accurate mapping of the yeast proteomic environment. Genome Biol., 9, R167.

40. Hořejší, Z., Takai, H., Adelman, C.A., Collis, S.J., Flynn, H., Maslen, S., Skehel, J.M., de Lange, T., Boulton, S.J., Horejsi, Z., et al. (2010) CK2 phospho-dependent binding of R2TP complex to TEL2 is essential for mTOR and SMG1 stability. Mol. Cell, 39, 839– 850.

41. Hořejší, Z., Stach, L., Flower, T.G., Joshi, D., Flynn, H., Skehel, J.M., O’Reilly, N.J., Ogrodowicz, R.W., Smerdon, S.J. and Boulton, S.J. (2014) Phosphorylation-Dependent PIH1D1 Interactions Define Substrate Specificity of the R2TP cochaperone complex. Cell Rep., 7, 19–26.

42. Pal, M., Morgan, M., Phelps, S.E.L., Roe, S.M., Parry-Morris, S., Downs, J.A., Polier, S., Pearl, L.H. and Prodromou, C. (2014) Structural basis for phosphorylation-dependent recruitment of Tel2 to Hsp90 by Pih1. Structure, 22, 805–818.

43. Ahmed, S., Alpi, A., Hengartner, M.O. and Gartner, A. (2001) C. elegans RAD-5/CLK-2 defines a new DNA damage checkpoint protein. Curr. Biol., 11, 1934–1944.

44. Shikata, M., Ishikawa, F. and Kanoh, J. (2007) Tel2 is required for activation of the Mrc1- mediated replication checkpoint. J. Biol. Chem., 282, 5346–5355.

45. Izumi, N., Yamashita, A., Hirano, H. and Ohno, S. (2012) Heat shock protein 90 regulates phosphatidylinositol 3-kinase-related protein kinase family proteins together with the RUVBL1/2 and Tel2-containing co-factor complex. Cancer Sci., 103, 50–57.

46. Kim, S.G., Hoffman, G.R., Poulogiannis, G., Buel, G.R., Jang, Y.J., Lee, K.W., Kim, B.-Y., Erikson, R.L., Cantley, L.C., Choo, A.Y., et al. (2013) Metabolic stress controls mTORC1 lysosomal localization and dimerization by regulating the TTT-RUVBL1/2 complex. Mol. Cell, 49, 172–85.

47. Rao, F., Cha, J., Xu, J., Xu, R., Vandiver, M.S., Tyagi, R., Tokhunts, R., Koldobskiy, M.A., Fu, C., Barrow, R., et al. (2014) Inositol Pyrophosphates Mediate the DNA-PK/ATM-p53 Cell Death Pathway by Regulating CK2 Phosphorylation of Tti1/Tel2. Mol. Cell, 54, 119–132.

48. David-Morrison, G., Xu, Z., Rui, Y.-N., Charng, W.-L., Jaiswal, M., Yamamoto, S., Xiong, B., Zhang, K., Sandoval, H., Duraine, L., et al. (2016) WAC Regulates mTOR Activity by Acting as an Adaptor for the TTT and Pontin/Reptin Complexes. Dev. Cell, 36, 139– 151.

49. Brown, M.C. and Gromeier, M. (2017) MNK Controls mTORC1:Substrate Association through Regulation of TELO2 Binding with mTORC1. Cell Rep., 18, 1444–1457.

50. Elías-Villalobos, A., Toullec, D., Faux, C., Séveno, M. and Helmlinger, D. (2019) Chaperone-mediated ordered assembly of the SAGA and NuA4 transcription co- activator complexes in yeast. Nat. Commun., 10, 5237.

51. Inoue, H., Sugimoto, S., Takeshita, Y., Takeuchi, M., Hatanaka, M., Nagao, K., Hayashi, T., Kokubu, A., Yanagida, M. and Kanoh, J. (2017) CK2 phospho-independent assembly of the Tel2-associated stress-signaling complexes in Schizosaccharomyces pombe. Genes to Cells, 22, 59–70.

52. Nishimura, K., Fukagawa, T., Takisawa, H., Kakimoto, T. and Kanemaki, M. (2009) An auxin-based degron system for the rapid depletion of proteins in nonplant cells. Nat. Methods, 6, 917–22.

53. Holland, A.J., Fachinetti, D., Han, J.S. and Cleveland, D.W. (2012) Inducible, reversible system for the rapid and complete degradation of proteins in mammalian cells. Proc. Natl. Acad. Sci. U. S. A., 109, E3350–7.

54. Hoke, S.M., Irina Mutiu, A., Genereaux, J., Kvas, S., Buck, M., Yu, M., Gloor, G.B. and Brandl, C.J. (2010) Mutational analysis of the C-terminal FATC domain of Saccharomyces cerevisiae Tra1. Curr. Genet., 56, 447–465.

55. Helmlinger, D., Marguerat, S., Villén, J., Swaney, D.L., Gygi, S.P., Bähler, J., Winston, F., Villen, J., Swaney, D.L., Gygi, S.P., et al. (2011) Tra1 has specific regulatory roles, rather than global functions, within the SAGA co-activator complex. EMBO J., 30, 2843–2852.

56. Herceg, Z. (2003) Genome-wide analysis of gene expression regulated by the HAT cofactor Trrap in conditional knockout cells. Nucleic Acids Res., 31.

57. Nagy, Z., Riss, A., Romier, C., le Guezennec, X., Dongre, A.R., Orpinell, M., Han, J., Stunnenberg, H. and Tora, L. (2009) The Human SPT20-Containing SAGA Complex Plays a Direct Role in the Regulation of Endoplasmic Reticulum Stress-Induced Genes. Mol. Cell. Biol., 29, 1649–1660.

58. Lang, G., Bonnet, J., Umlauf, D., Karmodiya, K., Koffler, J., Stierle, M., Devys, D. and Tora, L. (2011) The tightly controlled deubiquitination activity of the human SAGA complex differentially modifies distinct gene regulatory elements. Mol. Cell. Biol., 31, 3734–44.

59. Papai, G., Frechard, A., Kolesnikova, O., Crucifix, C., Schultz, P. and Ben-Shem, A. (2020) Structure of SAGA and mechanism of TBP deposition on gene promoters. Nature, 577, 711–716.

60. Wang, H., Dienemann, C., Stützer, A., Urlaub, H., Cheung, A.C.M. and Cramer, P. (2020) Structure of the transcription coactivator SAGA. Nature, 10.1038/s41586-020-1933-5.

61. Herbst, D.A., Esbin, M.N., Louder, R.K., Dugast-Darzacq, C., Dailey, G.M., Fang, Q., Darzacq, X., Tjian, R. and Nogales, E. (2021) Structure of the human SAGA coactivator complex: The divergent architecture of human SAGA allows modular coordination of transcription activation and co-transcriptional splicing. bioRxiv, 10.1101/2021.02.08.430339.

62. Jaenicke, L.A., von Eyss, B., Carstensen, A., Wolf, E., Xu, W., Greifenberg, A.K., Geyer, M., Eilers, M. and Popov, N. (2016) Ubiquitin-Dependent Turnover of MYC Antagonizes MYC/PAF1C Complex Accumulation to Drive Transcriptional Elongation. Mol. Cell, 61, 54–67.

63. Li, Y., Choi, P.S., Casey, S.C., Dill, D.L. and Felsher, D.W. (2014) MYC through miR-17-92 suppresses specific target genes to maintain survival, autonomous proliferation, and a Neoplastic state. Cancer Cell, 26, 262–272.

64. Subramanian, A., Tamayo, P., Mootha, V.K., Mukherjee, S., Ebert, B.L., Gillette, M.A., Paulovich, A., Pomeroy, S.L., Golub, T.R., Lander, E.S., et al. (2005) Gene set enrichment analysis: A knowledge-based approach for interpreting genome-wide expression profiles. Proc. Natl. Acad. Sci., 102, 15545–15550.

65. Liberzon, A., Birger, C., Thorvaldsdóttir, H., Ghandi, M., Mesirov, J.P. and Tamayo, P. (2015) The Molecular Signatures Database Hallmark Gene Set Collection. Cell Syst., 1, 417–425.

66. Schoggins, J.W. (2019) Interferon-Stimulated Genes: What Do They All Do? Annu. Rev. Virol., 6, 567–584.

67. Sathyan, K.M., McKenna, B.D., Anderson, W.D., Duarte, F.M., Core, L. and Guertin, M.J. (2019) An improved auxin-inducible degron system preserves native protein levels and enables rapid and specific protein depletion. Genes Dev., 33, 1441–1455.

68. Wang, W., Xu, L., Su, J., Peppelenbosch, M.P. and Pan, Q. (2017) Transcriptional Regulation of Antiviral Interferon-Stimulated Genes. Trends Microbiol., 25, 573–584.

69. Motwani, M., Pesiridis, S. and Fitzgerald, K.A. (2019) DNA sensing by the cGAS–STING pathway in health and disease. Nat. Rev. Genet., 10.1038/s41576-019-0151-1.

70. Negishi, H., Taniguchi, T. and Yanai, H. (2018) The interferon (IFN) class of cytokines and the IFN regulatory factor (IRF) transcription factor family. Cold Spring Harb. Perspect. Biol., 10, 1–16.

71. Field, A.K., Tytell, A.A., Lampson, G.P. and Hilleman, M.R. (1967) Inducers of interferon and host resistance. II. Multistranded synthetic polynucleotide complexes. Proc. Natl. Acad. Sci. U. S. A., 58, 1004–1010.

72. Kolosenko, I., Fryknäs, M., Forsberg, S., Johnsson, P., Cheon, H., Holvey-Bates, E.G., Edsbäcker, E., Pellegrini, P., Rassoolzadeh, H., Brnjic, S., et al. (2015) Cell crowding induces interferon regulatory factor 9, which confers resistance to chemotherapeutic drugs. Int. J. Cancer, 136, E51–E61.

73. Fink, K. and Grandvaux, N. (2013) STAT2 and IRF9. Jak-Stat, 2, e27521.

74. Cheon, H., Holvey-Bates, E.G., Schoggins, J.W., Forster, S., Hertzog, P., Imanaka, N., Rice, C.M., Jackson, M.W., Junk, D.J. and Stark, G.R. (2013) IFNβ-dependent increases in STAT1, STAT2, and IRF9 mediate resistance to viruses and DNA damage. EMBO J., 32, 2751–2763.

75. Sung, P.S., Cheon, H.J., Cho, C.H., Hong, S.H., Park, D.Y., Seo, H. Il, Park, S.H., Yoon, S.K., Stark, G.R. and Shin, E.C. (2015) Roles of unphosphorylated ISGF3 in HCV infection and interferon responsiveness. Proc. Natl. Acad. Sci. U. S. A., 112, 10443–10448.

76. Platanitis, E., Demiroz, D., Schneller, A., Fischer, K., Capelle, C., Hartl, M., Gossenreiter, T., Müller, M., Novatchkova, M. and Decker, T. (2019) A molecular switch from STAT2-IRF9 to ISGF3 underlies interferon-induced gene transcription. Nat. Commun., 10, 1–17.

77. Zambelli, F., Pesole, G. and Pavesi, G. (2009) Pscan: Finding over-represented transcription factor binding site motifs in sequences from co-regulated or co-expressed genes. Nucleic Acids Res., 37, 247–252.

78. Imrichová, H., Hulselmans, G., Atak, Z.K., Potier, D. and Aerts, S. (2015) I-cisTarget 2015 update: Generalized cis-regulatory enrichment analysis in human, mouse and fly. Nucleic Acids Res., 43, W57–W64.

79. Meers, M.P., Bryson, T.D., Henikoff, J.G. and Henikoff, S. (2019) Improved cut&run chromatin profiling tools. Elife, 8, 1–16.

80. Skene, P.J. and Henikoff, S. (2017) An efficient targeted nuclease strategy for high-resolution mapping of DNA binding sites. Elife, 6, 1–35.

81. Genereaux, J., Kvas, S., Dobransky, D., Karagiannis, J., Gloor, G.B. and Brandl, C.J. (2012) Genetic evidence links the ASTRA protein chaperone component Tti2 to the SAGA transcription factor Tra1. Genetics, 191, 765–780.

82. Antonova, S. V., Haffke, M., Corradini, E., Mikuciunas, M., Low, T.Y., Signor, L., van Es, R.M., Gupta, K., Scheer, E., Vos, H.R., et al. (2018) Chaperonin CCT checkpoint function in basal transcription factor TFIID assembly. Nat. Struct. Mol. Biol., 25, 1119– 1127.

83. Kassem, S., Villanyi, Z. and Collart, M.A. (2017) Not5-dependent co-translational assembly of Ada2 and Spt20 is essential for functional integrity of SAGA. Nucleic Acids Res., 45, 1186–1199.

84. Kamenova, I., Mukherjee, P., Conic, S., Mueller, F., El-Saafin, F., Bardot, P., Garnier, J.-M., Dembele, D., Capponi, S., Timmers, H.T.M.M., et al. (2019) Co-translational assembly of mammalian nuclear multisubunit complexes. Nat. Commun., 10, 1740.

85. Nikiforov, M.A., Chandriani, S., Park, J., Kotenko, I., Matheos, D., Johnsson, A., McMahon, S.B. and Cole, M.D. (2002) TRRAP-dependent and TRRAP-independent transcriptional activation by Myc family oncoproteins. Mol. Cell. Biol., 22, 5054–63.

86. Blaszczyk, K., Olejnik, A., Nowicka, H., Ozgyin, L., Chen, Y.-L., Chmielewski, S., Kostyrko, K., Wesoly, J., Balint, B.L., Lee, C.-K., et al. (2015) STAT2/IRF9 directs a prolonged ISGF3-like transcriptional response and antiviral activity in the absence of STAT1. Biochem. J., 466, 511–524.

87. Au-Yeung, N. and Horvath, C.M. (2018) Histone H2A.Z Suppression of Interferon-Stimulated Transcription and Antiviral Immunity is Modulated by GCN5 and BRD2. iScience, 6, 68–82.

88. Jin, Q., Zhuang, L., Lai, B., Wang, C., Li, W., Dolan, B., Lu, Y., Wang, Z., Zhao, K., Peng, W., et al. (2014) Gcn5 and PCAF negatively regulate interferon-β production through HAT- independent inhibition of TBK1. EMBO Rep., 15, 1192–201.

89. Rajagopalan, D., Tirado-Magallanes, R., Bhatia, S.S., Teo, W.S., Sian, S., Hora, S., Lee, K.K., Zhang, Y., Jadhav, S.P., Wu, Y., et al. (2018) TIP60 represses activation of endogenous retroviral elements. Nucleic Acids Res., 10.1093/nar/gky659.

90. Chiappinelli, K.B., Strissel, P.L., Desrichard, A., Li, H., Henke, C., Akman, B., Hein, A., Rote, N.S., Cope, L.M., Snyder, A., et al. (2015) Inhibiting DNA Methylation Causes an Interferon Response in Cancer via dsRNA Including Endogenous Retroviruses. Cell, 162, 974–986.

91. Roulois, D., Yau, H.L., Pugh, T.J., Brien, O., Carvalho, D. D. De, Roulois, D., Yau, H.L., Singhania, R., Wang, Y., Danesh, A., et al. (2015) DNA-Demethylating Agents Target Colorectal Cancer Cells by Inducing Viral Mimicry by Endogenous Transcripts. Cell, 162, 961–973.

92. Bailetti, A.A., Negrón-Pineiro, L.J., Dhruva, V., Harsh, S., Lu, S., Bosula, A. and Bach, E.A. (2019) Enhancer of Polycomb and the Tip60 complex repress hematological tumor initiation by negatively regulating JAK/STAT pathway activity. DMM Dis. Model. Mech., 12.

93. Wang, L. and Dent, S.Y. (2014) Functions of SAGA in development and disease. Epigenomics, 6, 329–339.

94. Judes, G., Rifaï, K., Ngollo, M., Daures, M., Bignon, Y.J., Penault-Llorca, F. and Bernard- Gallon, D. (2015) A bivalent role of TIP60 histone acetyl transferase in human cancer. Epigenomics, 7, 1351–1363.

95. Gongora, C., Candeil, L., Vezzio, N., Copois, V., Denis, V., Breil, C., Molina, F., Fraslon, C., Conseiller, E., Pau, B., et al. (2008) Altered expression of cell proliferation-related and interferon-stimulated genes in colon cancer cells resistant to SN38. Cancer Biol. Ther., 7, 822–832.

96. Khodarev, N.N., Beckett, M., Labay, E., Darga, T., Roizman, B. and Weichselbaum, R.R. (2004) STAT1 is overexpressed in tumors selected for radioresistance and confers protection from radiation in transduced sensitive cells. Proc. Natl. Acad. Sci., 101, 1714–1719.

97. Weichselbaum, R.R., Ishwaran, H., Yoon, T., Nuyten, D.S.A., Baker, S.W., Khodarev, N., Su, A.W., Shaikh, A.Y., Roach, P., Kreike, B., et al. (2008) An interferon-related gene signature for DNA damage resistance is a predictive marker for chemotherapy and radiation for breast cancer. Proc. Natl. Acad. Sci. U. S. A., 105, 18490–5.

98. Khodarev, N.N., Minn, A.J., Efimova, E. V., Darga, T.E., Labay, E., Beckett, M., Mauceri, H.J., Roizman, B. and Weichselbaum, R.R. (2007) Signal transducer and activator of transcription 1 regulates both cytotoxic and prosurvival functions in tumor cells. Cancer Res., 67, 9214–9220.

99. Fryknäs, M., Dhar, S., Öberg, F., Rickardson, L., Rydåker, M., Göransson, H., Gustafsson, M., Pettersson, U., Nygren, P., Larsson, R., et al. (2007) STAT1 signaling is associated with acquired crossresistance to doxorubicin and radiation in myeloma cell lines. Int. J. Cancer, 120, 189–195.

100. Suzuki, K., Bose, P., Leong-Quong, R.Y., Fujita, D.J. and Riabowol, K. (2010) REAP: A two minute cell fractionation method. BMC Res. Notes, 3, 294.

101. Schwalb, B., Michel, M., Zacher, B., Frühauf, K., Demel, C., Tresch, A., Gagneur, J. and Cramer, P. (2016) TT-seq maps the human transient transcriptome. Science (80-. )., 352, 1225–1228.

102. Kim, D., Langmead, B. and Salzberg, S.L. (2015) HISAT: A fast spliced aligner with low memory requirements. Nat. Methods, 12, 357–360.

103. Anders, S., Pyl, P.T. and Huber, W. (2015) HTSeq-A Python framework to work with high-throughput sequencing data. Bioinformatics, 31, 166–169.

104. Love, M.I., Huber, W. and Anders, S. (2014) Moderated estimation of fold change and dispersion for RNA-seq data with DESeq2. Genome Biol., 15, 1–21.

105. Afgan, E., Baker, D., Batut, B., Van Den Beek, M., Bouvier, D., Ech, M., Chilton, J., Clements, D., Coraor, N., Grüning, B.A., et al. (2018) The Galaxy platform for accessible, reproducible and collaborative biomedical analyses: 2018 update. Nucleic Acids Res., 46, W537–W544.

106. Cong, L., Ran, F.A., Cox, D., Lin, S., Barretto, R., Habib, N., Hsu, P.D., Wu, X., Jiang, W., Marraffini, L.A., et al. (2013) Multiplex Genome Engineering Using CRISPR/Cas Systems. Science (80-. )., 339, 819–823.

107. Lo, R. and Matthews, J. (2012) High-resolution genome-wide Mapping of AHR and ARNT binding sites by ChIP-Seq. Toxicol. Sci., 130, 349–361.

