## Supplementary material for "The TRRAP transcription cofactor represses interferon-stimulated genes in colorectal cancer cells": AllSupplementalFiguresandTables4-6

Detilleux D. *et al.*, Supplemental Figure 1

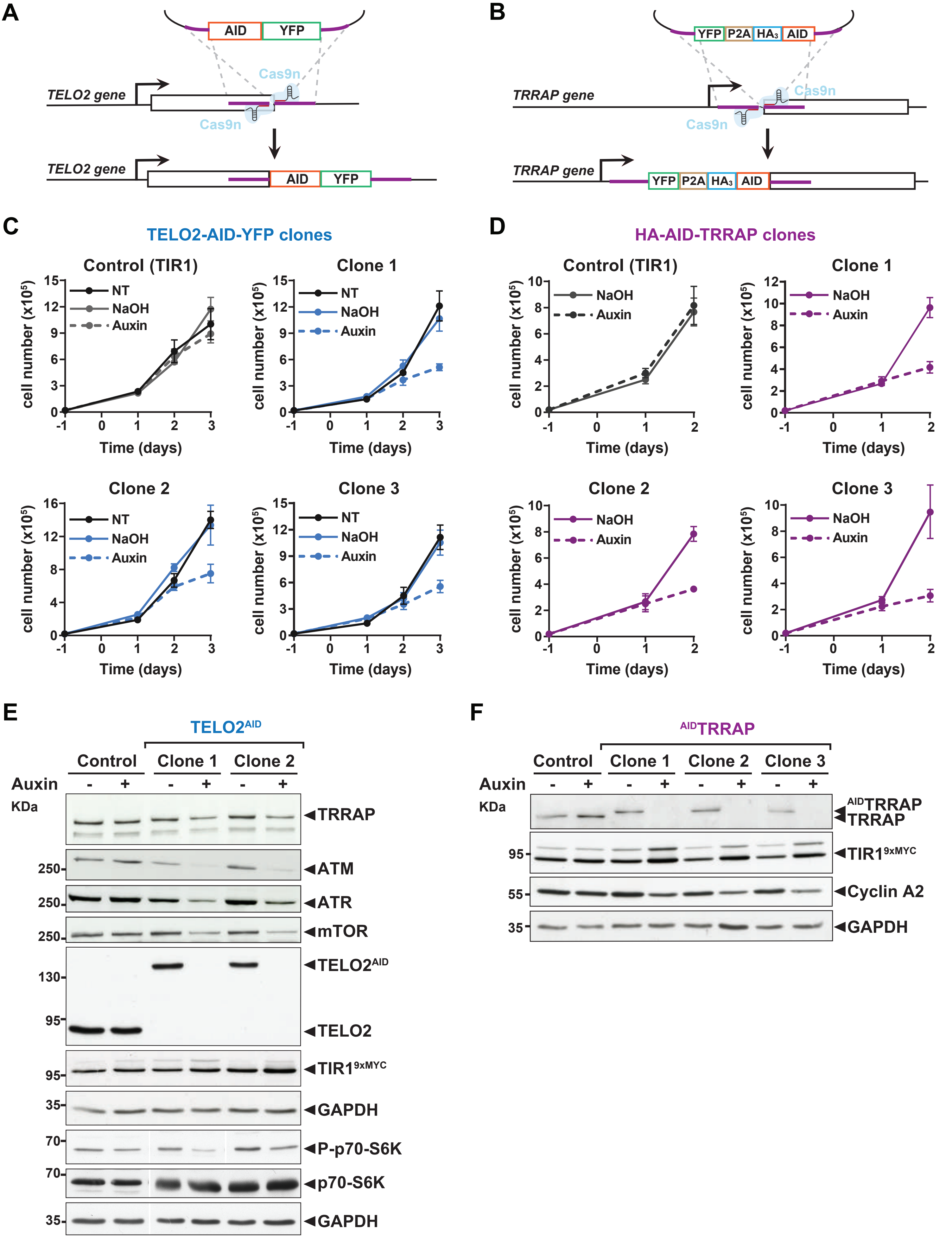

**Supplemental Figure 1: Characterization of TELO2-AID and AID-TRRAP cell lines.**

(A,B) Schematic depiction of the strategy designed to tag endogenous TELO2 (A) and TRRAP (B) with an auxin-inducible degron (AID) in in an HCT116 cell line stably expressing *Oryza sativa* TIR1. Gene editing was performed using the D10A nickase Cas9n mutant (105), which facilitates homology-directed repair and reduces off targets. Repair donor plasmids comprise sequences encoding a full-length IAA17-derived degron (50), a YFP fluorescent tag, and three HA epitopes. To reduce the risks of affecting TRRAP function with a long fusion sequence at its N-terminus, we inserted a 2A peptide (P2A) between HA-AID and YFP, which is then cleaved off during translation.

(C,D) Proliferation rates of parental HCT116-OsTIR1 cells (Control) and three distinct $TELO2^{AID}$ (C) or $^{AID}TRRAP$ (D) clones. The number of viable cells was measured using the trypan blue exclusion assay. 20,000 cells were plated at day -1. At day 0, cells were treated with auxin (dashed line), vehicle (NaOH, full line), or left untreated (NT, black) and monitored at the indicated time points.

(E,F) Immunoblotting of the indicated proteins in extracts from HCT116-OsTIR1 cells (Control), two distinct $TELO2^{AID}$ clones treated with auxin (+) or NaOH (-) for 48 hours (E), and three distinct $^{AID}TRRAP$ clones treated with auxin (+) or NaOH (-) for 24 hours. The OsTIR1 F-box protein was detected using an anti-MYC antibody. GAPDH was used as an internal control for equal loading.

**A**

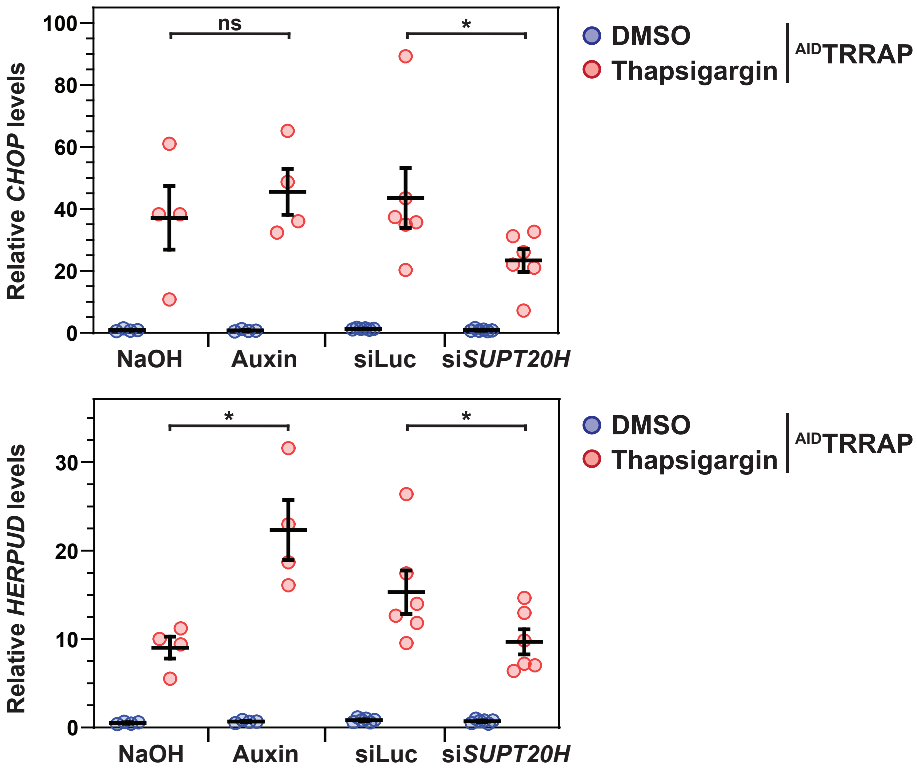
Relative *CHOP* levels
100
80
60
40
20
0
ns
*
NaOH
Auxin
siLuc
si*SUPT20H*
DMSO
Thapsigargin
AIDTRRAP
Relative *HERPUD* levels
30
20
10
0
*
*
NaOH
Auxin
siLuc
si*SUPT20H*
DMSO
Thapsigargin
AIDTRRAP

**B**

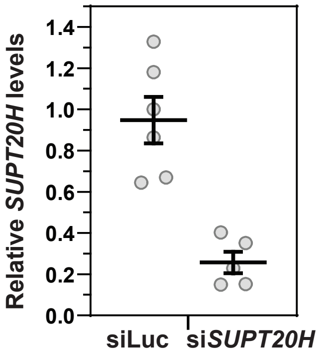
Relative *SUPT20H* levels
1.4
1.2
1.0
0.8
0.6
0.4
0.2
0.0
siLuc
si*SUPT20H*

**Supplemental Figure 2: TRRAP is not involved in SAGA-dependent ER stress gene induction.**

(A) Expression of *CHOP* and *HERPUD* using quantitative RT–PCR of RNA extracted from $^{AID}$TRRAP cells treated with either thapsigargin (red dots) or DMSO (blue dots) for 3 hours. Prior to ER stress induction, $^{AID}$TRRAP cells were either treated with auxin or NaOH for 48 hours, or transfected with siRNAs targeting Luciferase (Luc) or *SUPT20H* for 48 hours. *PPIB* served as a control for normalization across samples. Values from one control sample (DMSO, NaOH) were set to 1 to allow comparisons across conditions and replicates. Each line represents the mean value of either four (NaOH, Auxin) or six (siLuc, si*SUPT20H*) independent experiments, overlaid with individual data points and error bars showing the standard error of the mean (SEM). Statistical significance was determined by two-way ANOVA followed by Tukey's multiple comparison tests. *$P \leq 0.05$; ns: $P > 0.05$.

(B) Quantitative RT–PCR analysis of *SUPT20H* expression normalized to *PPIB* upon siRNA-mediated knockdown, as compared to control siRNAs targeting Luciferase. RNAs were from $^{AID}$TRRAP cells 48 hours after transfection. Each line represents the mean value of six independent experiments, overlaid with individual data points and error bars showing the SEM.

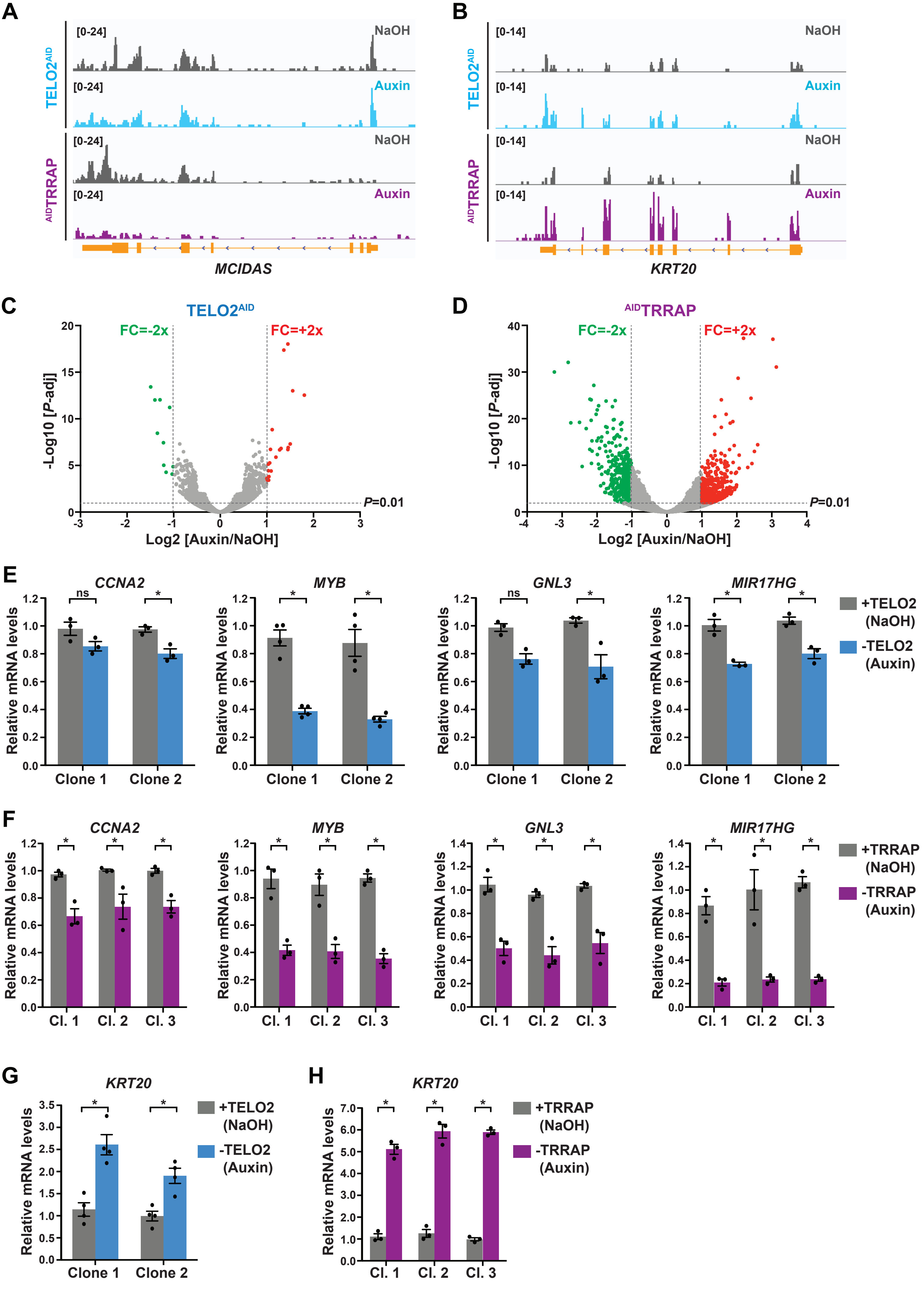
A
[0-24]
NaOH
TELO2AID
[0-24]
Auxin
AIDTRRAP
[0-24]
NaOH
[0-24]
Auxin
MCIDAS
B
[0-14]
NaOH
[0-14]
Auxin
[0-14]
NaOH
[0-14]
Auxin
KRT20
C
TELO2AID
FC=-2x
FC=+2x
-Log10 [P-adj]
P=0.01
Log2 [Auxin/NaOH]
D
AIDTRRAP
E
CCNA2
MYB
GNL3
MIR17HG
Relative mRNA levels
ns
*
Clone 1
Clone 2
+TELO2 (NaOH)
-TELO2 (Auxin)
F
Cl. 1
Cl. 2
Cl. 3
+TRRAP (NaOH)
-TRRAP (Auxin)
G
KRT20
H

**Supplemental Figure 3: Gene expression changes upon TELO2 and TRRAP depletion.**

(A,B) Scaled IGV (Integrative genomics viewer) snapshots of two differentially expressed genes, *MCIDAS* (A) and *KRT20* (B) from RNA sequencing (RNA-seq) performed in three distinct $TELO2^{AID}$ and $^{AID}TRRAP$ clones treated with auxin or NaOH for 48h or 24h, respectively.

(C,D) Volcano plots showing transcript level fold change (FC) measured upon TELO2 (C) and TRRAP (D) depletion plotted against the adjusted *P* value (FDR). FC were calculated as the Log2 of the ratio of the expression value of each gene between NaOH- (+TELO2 or +TRRAP) and auxin-treated (-TELO2 or -TRRAP) cells. Two-fold change thresholds and a 1% FDR cut-off are shown. Blue dots represent mRNAs which levels decrease at least 2-fold and red dots represent mRNAs which levels increase at least 2-fold.

(E-H) Quantitative RT-PCR analysis of selected downregulated (*CCNA2*, *MYB*, *GNL3*, and *MIR17HG*) (E,F) and one upregulated gene (*KRT20*) (G,H). mRNAs levels were measured in two $TELO2^{AID}$ clones treated with auxin or NaOH for 48h (E,G), and in three $^{AID}TRRAP$ clones treated with auxin or NaOH for 24h (F,H). Each value represents mean mRNA levels from at least three independent experiments, overlaid with individual data points and error bars showing the SD. *PPIB* served as a control for normalization across samples. Values from one NaOH-treated control sample were set to 1, allowing comparisons across culture conditions and replicates. Statistical significance was determined by two-way ANOVA followed by Bonferroni's multiple comparison tests. *P ≤ 0.01.

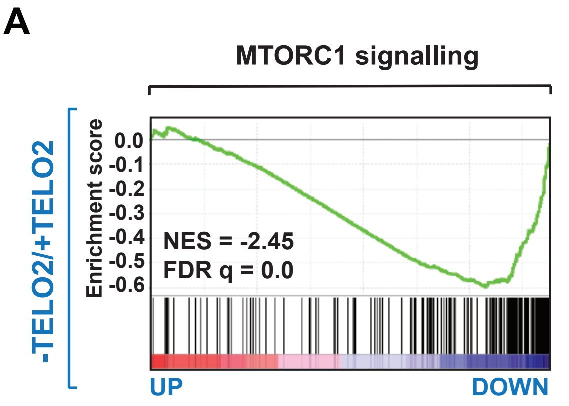
A
MTORC1 signalling
-TELO2/+TELO2
Enrichment score
0.0
-0.1
-0.2
-0.3
-0.4
-0.5
-0.6
NES = -2.45
FDR q = 0.0
UP
DOWN

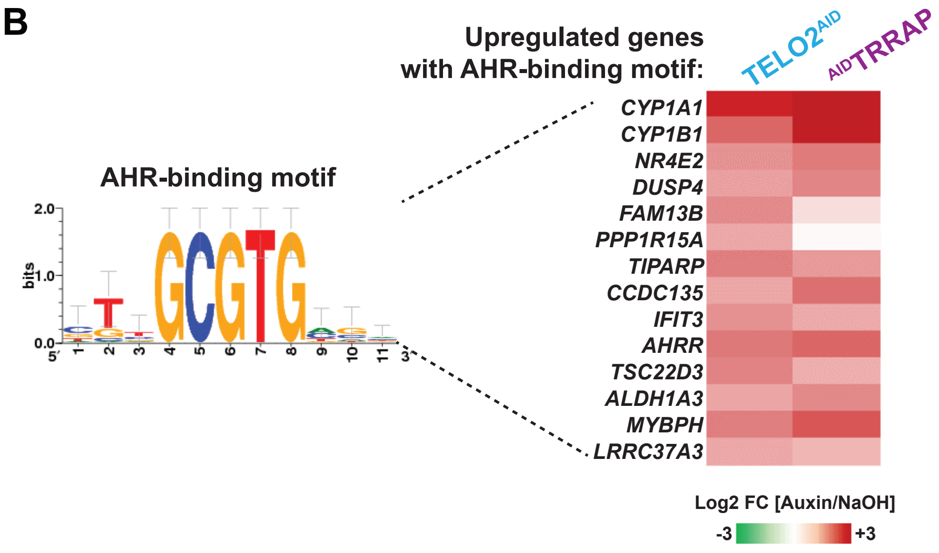
B
AHR-binding motif
bits
2.0
1.0
0.0
5'
1
2
3
4
5
6
7
8
9
10
11
3'
Upregulated genes
with AHR-binding motif:
TELO2AID
AIDTRRAP
CYP1A1
CYP1B1
NR4E2
DUSP4
FAM13B
PPP1R15A
TIPARP
CCDC135
IFIT3
AHRR
TSC22D3
ALDH1A3
MYBPH
LRRC37A3
Log2 FC [Auxin/NaOH]
-3
+3

**Supplemental Figure 4: Auxin treatment induces AHR-responsive genes.**

(A) Gene set enrichment analysis (GSEA) showing the enrichment of genes encoding MTORC1 signaling components in the ranked transcriptome profiles of TELO2-depleted cells. Green lines represent enrichment profiles. NES: normalized enrichment score. Each hit from the hallmark gene set is represented by a vertical black bar, positioned on the ranked transcriptome profile with color-coded fold change values.

(B) Heat map representation of genes containing a consensus AHR binding motif and which expression changes in TELO2 and TRRAP-depleted cells. The Log2 ratio between auxin- and NaOH-treated cells for each transcript is represented using a sequential color scale. All data are from RNA-seq experiments performed in three distinct $TELO2^{AID}$ and $^{AID}TRRAP$ clones treated with auxin or NaOH for 48h or 24h, respectively. Shown is the sequence logo of the minimal core AHR-binding motif 5′-GCGTG-3′ (transfac_pro__M00778) (106).

Detilleux D. *et al.*, Supplemental Figure 5

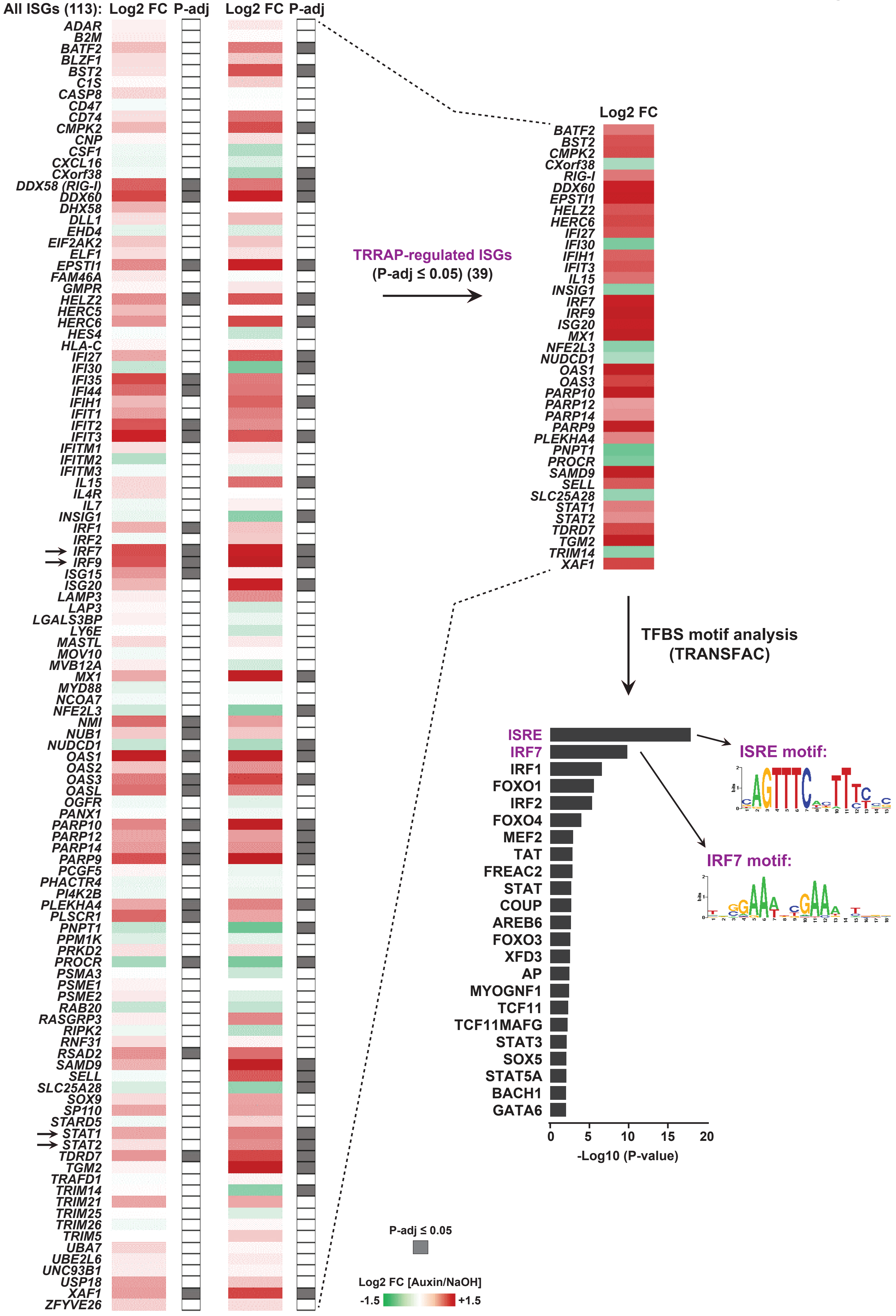

**Supplemental Figure 5: TELO2 and TRRAP repress type I interferon stimulated genes.**

Heat map representation of the expression of a list of IFN-I stimulated genes (113 ISGs) in TELO2 and TRRAP-depleted cells. The list of 113 ISGs results from merging the list of 97 genes from the IFN $\alpha$-responsive hallmark set with the lists of genes regulated by U-ISGF3 (29) and ISGF3 (49), as defined in (73). The Log2 ratio between auxin- and NaOH-treated cells for each transcript is represented using a sequential color scale. Gray: adjusted P-values ≤ 0.05 (FDR ≤ 5%); white: > 0.05. All data are from RNA-seq experiments performed in three distinct $TELO2^{AID}$ and $^{AID}TRRAP$ clones treated with auxin or NaOH for 48h and 24h, respectively. Arrows indicate the genes encoding the IRF7 and ISGF3 transcription factors. A heat map highlighting the 39 ISGs which expression changes upon TRRAP depletion (FDR ≤ 5%) is shown on the right. Below is a histogram of *P*-values of transcription factor binding sites (TFBS) enrichment in the sequences of the 39 TRRAP-regulated ISGs. Shown are the sequence logos of the two most enriched motifs, the ISRE motif (transfac_public__M00258) and the IRF7 motif (transfac_public__M00453). The number of genes in each category is indicated in parentheses.

**Supplemental Table 4. List of siRNA sequences used in this study.**

| mRNA targeted | Sequence |
|---|---|
| *Firefly Luciferase (FFL)* | CUUACGCUGAGUACUUCGA |
| *ATXN7L3* | TGAATAAGTCTGAGAGTGA |
| *TIP60* | CGUCUGGAUGAAUGGGUGA |
| *p400* | UGAAGAAGGUUCCCAAGAA |
| *TADA3* | GCUCUGAGGAUGAUGGCAU |
| *TRRAP* | GCCCUGUUCUUUCGCUUUGUA |
| *SUPT20H* | UGAAGUAGAUGUGGAGAAA |

**Supplemental Table 5. List of oligonucleotides used in this study.**

| Gene | Name | Purpose | Region targeted | Sequence |
|---|---|---|---|---|
| *TELO2* | DHO 1067 | CRISPR-KI C-ter TELO2 sgRNA A | 3 prime neat stop codon | CACCGACCACCCTCGCCGACAGCA |
| *TELO2* | DHO 1068 | CRISPR-KI C-ter TELO2 sgRNA A | 3 prime near stop codon | AAACTGCTGTCGGCGAGGGTGGTC |
| *TELO2* | DHO 1069 | CRISPR-KI C-ter TELO2 sgRNA B | 3 prime near stop codon | CACCGAGGGACTAGGGAGACGCGGG |
| *TELO2* | DHO 1070 | CRISPR-KI C-ter TELO2 sgRNA B | 3 prime near stop codon | AAACCCCGCGTCTCCCTAGTCCCTC |
| *TRRAP* | DHO 1227 | CRISPR-KI N-ter TRRAP sgRNA C | 3 prime near start codon | CACCGGCGTTTGTTGCAACACAGG |
| *TRRAP* | DHO 1228 | CRISPR-KI N-ter TRRAP sgRNA C | 3 prime near start codon | AAACCCTGTGTTGCAACAAACGCC |
| *TRRAP* | DHO 1229 | CRISPR-KI N-ter TRRAP sgRNA D | 3 prime near start codon | CACCGCTTCTCAAGAGAAAAGTATC |
| *TRRAP* | DHO 1230 | CRISPR-KI N-ter TRRAP sgRNA D | 3 prime near start codon | AAACGATACTTTTCTCTTGAGAAGC |
| *PPIB* | DHO 1083 | RT-qPCR | | CTTCCCCGATGAGAACTTCAAACT |
| *PPIB* | DHO 1084 | RT-qPCR | | CACCTCCATGCCCTCTAGAACTTT |
| *CCNA2* | DHO 1146 | RT-qPCR | | AGATGAAAAGCCAGTGAGTGT |
| *CCNA2* | DHO 1147 | RT-qPCR | | GTGATGTCTGGCTGTTTCTTCA |
| *KRT20* | DHO 1114 | RT-qPCR | | TCATGGCCCAGAAGAACCTT |
| *KRT20* | DHO 1115 | RT-qPCR | | TTCACTGTGACCTGTTGCTG |
| *OAS1* | DHO 1389 | RT-qPCR | | AGTTGACTGGCGGCTATAAAC |
| *OAS1* | DHO 1390 | RT-qPCR | | GTGCTTGACTAGGCGGATGAG |
| *IFIT1* | DHO 1402 | RT-qPCR | | TGACGTCAATGCAATTATCCA |
| *IFIT1* | DHO 1403 | RT-qPCR | | GCCCGCTCATAGTACTCCAG |
| *IRF7* | DHO 1404 | RT-qPCR | | CGAGCTGCACGTTCCTATAC |
| *IRF7* | DHO 1405 | RT-qPCR | | AGCAGTTCCTCCGTGTAGC |
| *IRF9* | DHO 1497 | RT-qPCR | | TTCAGACATTGGGAGCAGCA |
| *IRF9* | DHO 1498 | RT-qPCR | | AGGAAGCAGAAACTCCAGGG |
| *ATXN7L3* | DHO 1541 | RT-qPCR | | AGATATACGCGGACCTGGTC |
| *ATXN7L3* | DHO 1542 | RT-qPCR | | AATTGGGGCAAACACACTCC |
| *RIG-I* | DHO 1578 | RT-qPCR | | GAGGATCTTCAGGCCCACT |
| *RIG-I* | DHO 1579 | RT-qPCR | | CTTGCTCCAGTTCCTCCAGA |
| *MYB* | DHO 1681 | RT-qPCR | | TGATGAAGACCCTGAGAAGGA |
| *MYB* | DHO 1682 | RT-qPCR | | GGTGTTCTCCCAAACAGGAA |
| *GNL3* | DHO 1685 | RT-qPCR | | GGGCTTTGCAAAACTGAGAA |
| *GNL3* | DHO 1686 | RT-qPCR | | TGGCCTCTTCTACCTGAGGA |
| *MIR17HG* | DHO 1691 | RT-qPCR | | GCCCAATCAAACTGTCCTGT |
| *MIR17HG* | DHO 1692 | RT-qPCR | | ACCGATCCCAACCTGTGTAG |
| *MX1* | DHO 1733 | RT-qPCR | | CGACACGAGTTCCACAAATG |
| *MX1* | DHO 1734 | RT-qPCR | | TGCCTTGATTTGCTGTTTCA |
| *KAT5* | DHO 1941 | RT-qPCR | | AATGTGGCCTGCATCCTAAC |
| *KAT5* | DHO 1942 | RT-qPCR | | TGTTTTCCCTTCCACTTTGG |
| *HERPUD* | DHO 1077 | RT-qPCR | | GGTTGGGGTCTTCAGTTTCAGG |
| *HERPUD* | DHO 1078 | RT-qPCR | | CTACTCCTCCCTGAGCAGATTC |
| *CHOP* | DHO 1101 | RT-qPCR | | AAGGCACTGAGCGTATCATGT |
| *CHOP* | DHO 1102 | RT-qPCR | | TGAAGATACACTTCCTTCTTGAAC |
| *SUPT20H* | DHO 1093 | RT-qPCR | | GCAGTCACCTGCACTGCAAACA |
| *SUPT20H* | DHO 1094 | RT-qPCR | | TCAGCTGAGGAGGAGGTGGACA |
| *PPIB-intron1* | DHO 1968 | nascent RT-qPCR | intron 1 | TCTCTCTCCCATCCTCAGGT |
| *PPIB-intron1* | DHO 1969 | nascent RT-qPCR | intron 1 | AGAGACCAAAGATCACCCGG |
| *IRF9-intron2* | DHO 1972 | nascent RT-qPCR | intron 2 | GTGGATCACGAGGTCAAGAGA |
| *IRF9-intron2* | DHO 1973 | nascent RT-qPCR | intron 2 | TCCCTGGTTCAAGCGATTCT |
| *OAS1-intron1* | DHO 1974 | nascent RT-qPCR | intron 1 | TGCAATGCCTTCAGAACAGT |
| *OAS1-exon2* | DHO 1975 | nascent RT-qPCR | exon 2 | AGTGGTGAGAGGACTGAGGA |
| *IRF7-intron2* | DHO 2000 | nascent RT-qPCR | intron 2 | GGGGAACATTCTCTGGGTCA |
| *IRF7-intron2* | DHO 2001 | nascent RT-qPCR | intron 2 | TCCTGGCTGTGAACCCTTAG |
| *MIR17HG* | DHO 1691 | CUTnRUN | gene body | GCCCAATCAAACTGTCCTGT |
| *MIR17HG* | DHO 1692 | CUTnRUN | gene body | ACCGATCCCAACCTGTGTAG |
| *MIR17HG* | DHO 1721 | CUTnRUN | promoter | CTTTGCAGTCTCGGGTGTTC |
| *MIR17HG* | DHO 1722 | CUTnRUN | promoter | CAACTGCTGTGGTCCTGATG |
| *IRF9* | DHO 1793 | CUTnRUN | promoter | GGCTAGAAAGTCCAGCCAGA |
| *IRF9* | DHO 1794 | CUTnRUN | promoter | GGCCCTTGTCAGTCTCACTC |
| *IRF7* | DHO 1945 | CUTnRUN | promoter | CCAGCTCTTGGCTCTACCC |
| *IRF7* | DHO 1946 | CUTnRUN | promoter | GCTCTGGCACCCAGGTACT |

**Supplemental Table 6. List of antibodies used in this study.**

| Protein targeted | Provider | Catalog number |
|---|---|---|
| ATR | CST | 13934 |
| ATM | CST | 2873 |
| SUPT7L | Bethyl | A-302-803A-1 |
| TRRAP | L. TORA lab | 2TRR-1B3 16 1 |
| TADA3 | L. TORA lab | 2678 |
| TELO2 | ProteinTech | 15975-1-AP |
| TTI1 | Santa Cruz Biotechnology | sc-271638 |
| GAPDH | Santa Cruz Biotechnology | sc-25778 |
| mTOR | CST | 2972 |
| phospho-p70 S6 (T389) | CST | 9234 |
| p70 S6 | CST | 9202 |
| TRRAP | L. TORA lab | 2TRR-2DR5 16 1 |
| SGF29 | L. TORA lab | 2461 |
| SUPT3H | L. TORA lab | 3118 |
| SUPT3H | L. TORA lab | 1-SP-2C10 |
| TAF10 | L. TORA lab | 2B11 |
| TADA1 | Novusbio | NBP1-56629 |
| IRF7 | Biolegend | 656002 |
| IRF3 | CST | 4302 |
| Phospho-IRF3 (S396) | Abcam | ab76293 |
| LaminA/C | CST | 2032 |
| RIG-I | CST | 3743 |
| RIG-I | Milipore | MABF297 |
| LamiA/C | CST | 4777 |
| MDA5 | CST | 5321 |
| MAVS | CST | 3993 |
| cMYC | CST | 13987 |
| IRF9 | Biolegend | 660702 |
| HA | Abcam | ab9110 |
| Tubulin | Sigma | T6074 |
| H2A.Z | Abcam | ab4174 |
